# Mechanistic Dissection of Entropic Penalty upon Ligand Binding and Molecular Flexibility via Molecular Dynamics Simulations and Machine Learning

**DOI:** 10.64898/2026.08.18.745526

**Authors:** Ta I Hung, Emily Vig, Chia-en Chang

## Abstract

Molecular flexibility governs how molecules behave, reorganize, and respond to their environment. Although experiments measure molar entropy for small molecules and molecular dynamics (MD) simulations capture molecular motions, quantifying configuration entropy and the concerted internal motions such as torsion rotations, angle bending, and their couplings are central to understanding thermodynamic behavior but remains challenging. To dissect these contributions, we used MD trajectories and developed an internal-coordinate PC-entropy (iPC-entropy) method to probe the origins of entropy and reveal how specific motions shape the thermodynamic landscape. The studies accurately captured molar entropy, identified key torsional motions as major contributors, and uncovered a critical angle–torsion coupling in which angle bending was strongly correlated with torsional rotation, a coupling that increases nonlinearly with molecular size. Evaluating entropic changes upon protein–ligand binding reveals that dominant entropic penalty arises from ligand dihedral rigidification rather than protein reorganization and highlights the specific dihedral rotations that become restricted. We also suggest systematic corrections for approaches considering solely rotamers to reliably reproduce the relative entropic penalty in computer-aided drug discovery. Together, our findings elucidate the molecular origins of entropy and entropy changes. In addition, we can quantify and illustrate the internal motions that strongly shape binding thermodynamics, thereby offering mechanistic insights to guide drug development.

## Introduction

Molecular motion plays a fundamental role in determining molecular properties, driving chemical reaction mechanisms and influencing the kinetics of biological processes ^1,2^. It also offers a critical pathway for understanding thermodynamics. Molecular motion provides valuable insights into how molecules behave and interact, although its complexity presents significant challenges. The flexibility of molecules allows them to adopt various configurations, which can be quantified by configurational entropy calculations. In essence, configurational entropy quantifies the disorder of molecules by measuring the number of possible configurations or states a molecule can adopt, which is essential for determining stability, reactivity, and behavior under various environments.

Configurational entropy is one of the most challenging thermodynamic properties to estimate because one must sample all possible configurations thoroughly. From a thermodynamic standpoint, molecules rotate, translate and vibrate to seek more configurations to maximize the entropy, which results in an energetically favorable state. Whether via its molecular recognition, the formation of a protein, crystal lattice, polymers and behavior of amorphous solid ^3–8^, the entropy contributes substantially to the driving force of molecular motion across both biological and non-biological molecules. A method to accurately dissect molecular motion and calculate the configurational entropy would provide valuable insights to understand molecular motion and the dynamic properties of molecules.

Entropy can be measured experimentally with calorimetry by measuring heats of fusion, transition and heat capacities over temperature ranges ^9^. However, as the size of the molecules increases, detecting the small heat changes becomes highly challenging and inaccurate. Recently, nuclear magnetic resonance (NMR) has provided valuable insights into estimating configurational entropy changes in biological systems ^10–13^. This approach provides clues to molecular motion and conformational transition. NMR can be used to study the effect of solvent interaction of molecular conformation. However, NMR requires extensive sampling of molecular motions and conformations, which can be time-consuming and is often limited to capturing only a subset of relevant motions but not the overall protein dynamics. Indeed, all experimental methods provide few or only single static structures without giving information on long-timescale correlated motion, which is important in understanding the mechanism and motion of small molecules and proteins^14^. Additionally, isotopically labeling small-molecule ligands and drugs are typically difficult and expensive ^10^. Computational methods are a complementary technique to bridge the knowledge gaps.

Computational methods have been widely used to quantify configurational entropy in chemical and biological systems, providing atomistic insight into how molecular flexibility contributes to molar entropy and binding affinity. Early approaches involved quasiharmonic (QH) approximation for computing configurational entropy directly from molecular dynamics (MD) simulation. QH approximation provides an analytical solution of configurational entropy by using multivariable Gaussian distribution to approximate the probability distribution function (PDF) of the internal coordinates of molecules ^15,16^. However, the QH approach assumes configurational PDF as a Gaussian distribution, which cannot correctly describe a real molecular system with multipeak and non-Gaussian PDF, thereby leading to over- or underestimating the configurational entropy ^17^. Other simulation-based methods such as histogramming and the use of torsion angle distribution compute entropies by estimating the probability distribution function of each state ^18,19^. Nearest-neighbor estimated configuration entropy ^20^, covariance analysis of dihedral angle in complex plan ^21^, and computing dihedral entropy of all rotatable bonds with Machine Learning ^22^ have been investigated. Non-simulation base methods such as the second-generation mining minima algorithm (M2) have provided insights into changes in configurational entropy upon binding for small-molecule HIV protease (HIVp) systems ^23^. M2 samples the configuration of molecules directly by searching all local minimums and directly obtaining the configurational integral ^24^. A thorough search is required to sample all molecular configurations, which can be computationally expensive.

Here, we post-processed MD trajectories to compute the molar entropy of organic molecules and the configurational entropy changes associated with protein–ligand binding (**Figure 1**). Using multi-fragmentation bond–angle–torsion (BAT) internal coordinates with angle-aware arithmetic to accurately represent molecular motions, an essential requirement for reliable entropy estimation, our approach applies unsupervised machine learning algorithm, principal component analysis (PCA), to decouple the correlated motions sampled in MD simulations into independent PC modes. We then used numerical integration of the Gibbs entropy formula to compute the configurational entropy. This internal-coordinate PC-entropy (iPC-entropy) method allows for directly calculating configurational entropy from MD data and identifies the dominant contributors to entropy. We analyzed and quantified the features contributing to molecular flexibility for each organic molecule as well as the entropy loss when the drugs ritonavir (RIT) and amprenavir (AMP) bind to their target HIVp. We also examined commonly used rotamer-counting approaches. The results highlight the sources of entropy and entropy loss upon protein–ligand binding, underscore the importance of system-dependent angle–torsion coupling, and may help formulate empirical models to improve entropy estimation for ligand screening and binding prediction.

**Figure 1.**
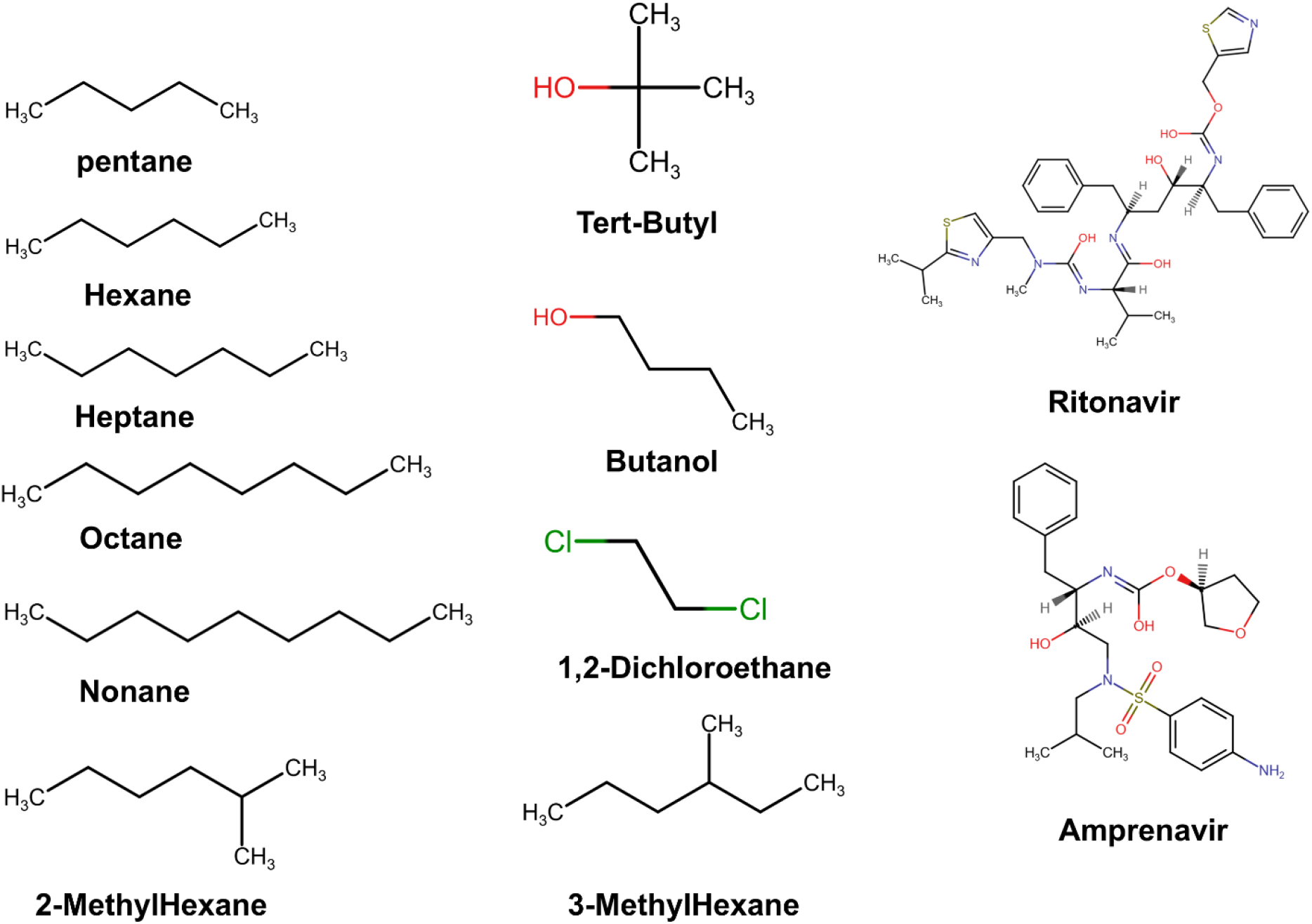
Organic molecules and HIV protease inhibitors ritonavir (RIT) and amprenavir (AMP).

### Theory

#### Molar entropy (S)

In general, molar entropy can be defined using Gibbs entropy formula:

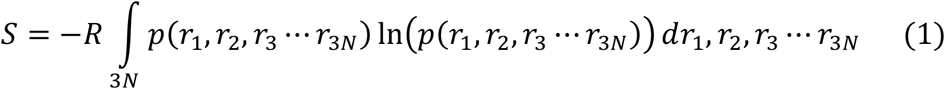

where *p*(*r*_1_, *r*_2_, *r*_3_ ⋯ *r*_3*N*_) is the probability distribution function (PDF) of all degrees of freedom (3N), and *r*_1_, *r*_2_, *r*_3_ ⋯ *r*_3*N*_ are cartesian coordinates of the atoms. N is the number of atoms.

External translational and rotational motions are independent with internal conformation. Therefore, Molar entropy (S) can be further separated into external term (S_ext_) which includes translational and rotational entropies (S_trans_ and S_rot_), and configurational entropy (S_config_).

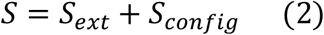

The S_ext_ accounts for 6 translational and rotational degrees of freedom and can be solved analytically.

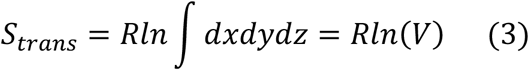

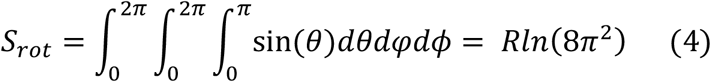

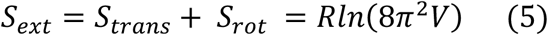

Therefore, the molar entropy (S) is derived as:

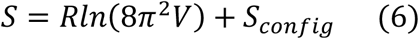

The integral of the external term can be solved analytically, and we only need to focus on the configurational term (*S*_*config*_) which can be written as:

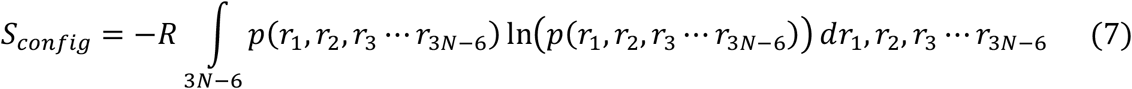

#### Bond-Angle-Torsions (BAT) Coordinate

To accurately describe molecular motions modeled by molecular mechanics force field, internal BAT coordinates are used. Classical BAT coordinates, also termed as Z-matrix, are defined as sets of root atoms and the position of other atoms depends on the previously defined atoms bonded in sequence. As illustrated in **Figure 2A**, Atom 1 is at the origin; atom 2 is defined by a bond length (b2) and placed on the z-axis; and Atom 3 is defined by a bond length (b3), bond angle (a3) and placed on the x–z plane, and thus we eliminate 6 degrees of freedoms. Total degree of freedom of molecules with N atoms has 3N-6 internal coordinates. Atoms 1–3 are termed root atoms for this molecule. Atoms i > 3 are defined by (bi, ai, ϕi), where t are torsion angles. For example, atom 4 is presented by (b4, a4, ϕ4). The internal coordinate is then constructed in tree. Starting from root atoms, all subsequent atoms are defined sequentially.

**Figure 2.**
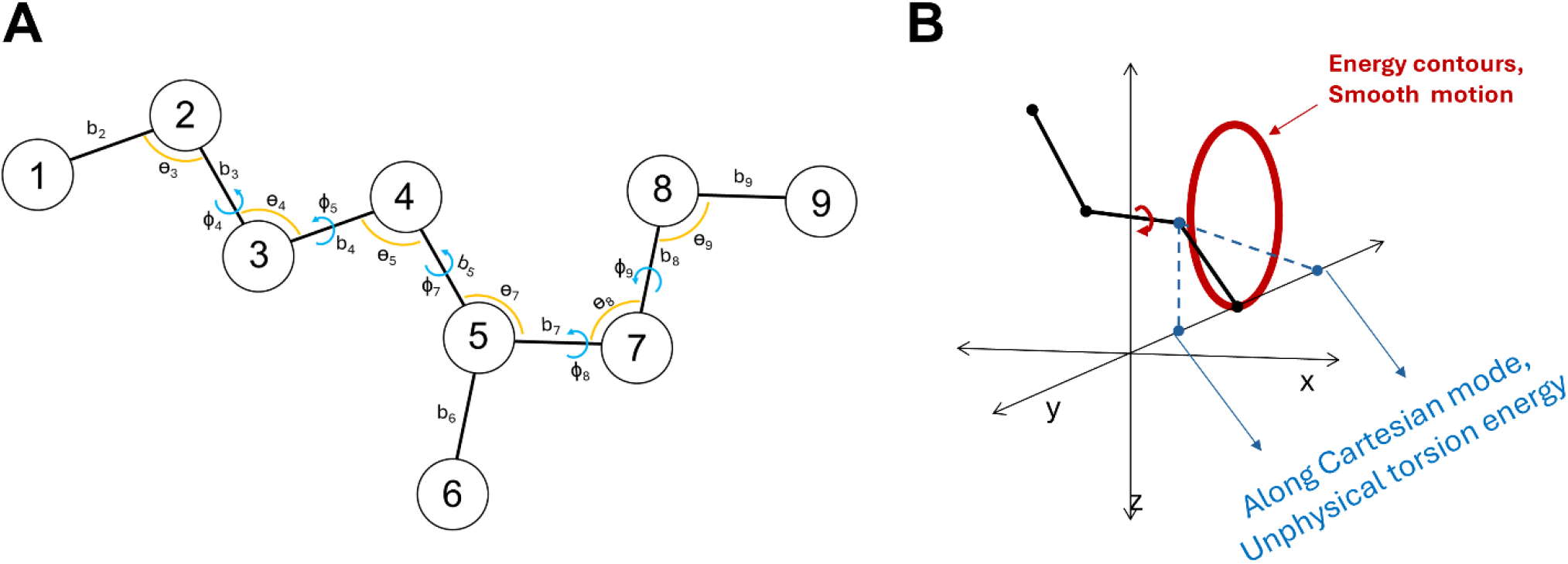
Describing molecular configuration using Bond-Angle-Torsion (BAT) internal Coordinate. **(A)** Classical internal BAT coordinate representation (Z-matrix) for a small molecule. Atoms 1−3 are termed root atoms for this molecule. Atoms i > 3 are defined by (bi, θi, ϕi), where b, a and θ are bond, angle and torsions, respectively. For example, atom 4 is presented by (b4, θ4, ϕ4). **(B)** Rotation of torsion angles follows a smooth circular line shown in red. Describing rotation using cartesian coordinates reduces circular motion to linear combination of Cartesian components. The projection will result in linear displacement in the y-axis which can stretch the bond and create unphysically high energy.

To accurately perform mathematical operations in BAT internal coordinates, we needed to address the discontinuity issue at (±180°) for torsion rotation. For instance, dihedral rotation from 179° to −179° is actually a 2° shift in angular space. However, the basic arithmetic operation will result in a large rotation, 179°-(−179°) = 358° ^25^. To address the discontinuity issue, we introduced angle-aware arithmetic by mapping each torsion angle θ on the unit circle: (cos θ, sin θ). In case of iPCA, we need to correctly compute angle-aware covariance matrix by using average, subtraction and addition in (cos θ, sin θ) space. For dihedral angles x and y with sample size N, covariance is calculated as:

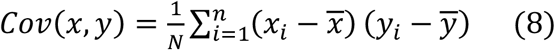

The angle-aware average value of the variable x is:

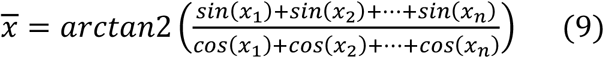

while angle-aware subtraction is:

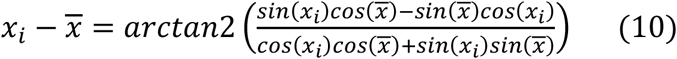

Using angle-aware average and subtraction, we can correctly compute the covariance matrix followed by the standard calculation of eigenvectors and eigenvalues. All conformations are then projected along each eigenvector. Similarly, projection is computed using the angle-aware mean and addition (dot product):

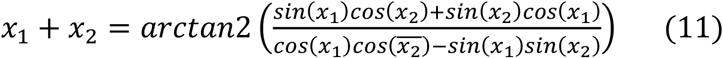

#### Multi-fragmentation BAT

In classical BAT, all atoms are defined in sequence, and thus it could encounter severe error propagation when defining large molecules such as proteins. A small artifact close to the root atoms will result in a large and unrealistic movement near the end of the protein. Instead of defining all atoms sequentially, we employed the multi-fragments approach to avoid error propagation ^26^. We fragment the proteins into N fragments based on the local structures such as alpha helix, beta sheet and flexible loops. Pseudo-bond from the first atom of the fragment to the root atoms is created to connect the fragments to root atoms. This approach allows us to correctly define the position of each fragment directly from the root atoms. Moreover, multi-fragmentation allows us to efficiently define protein dimer, protein-ligand complex and protein-protein complexes by simple pseudo-bond connection. In our case, HIVp is a dimeric protein, and thus we treat one of the subunits as fragment and connect the first atom to the root (**Figure S1A**).

#### Coordinate Transformation

Molecular configurations are described with commonly used Cartesian Coordinate *r* ≡ (*x, y, z*). The probability density function (PDF) can be expressed as *p(r)* and *S*_*config*_ = *S*_*r*_ = −*R* ∫_*configurration*_ *p*(*r*) ln(*p*(*r*)) *dr*. Other coordinates, such as the BAT coordinate *q* ≡ (*b, θ, ϕ*) introduced above can also be used to describe molecular configuration with Jacobians *J(q)*, where the PDF can be expressed by *q(r)*. Note that the integral of the PDF equals 1, regardless of the coordinate system used.

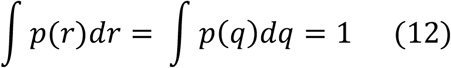

We know that *dr* = *J*(*q*)*dq*, plug it back to **equation (12)**, we obtained:

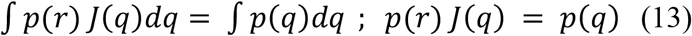

After coordinate transformation from coordinate *r* to *q*, PDF *p*(*q*) can be used to describe Gibbs entropy using coordinate *q*, S_*q*_:

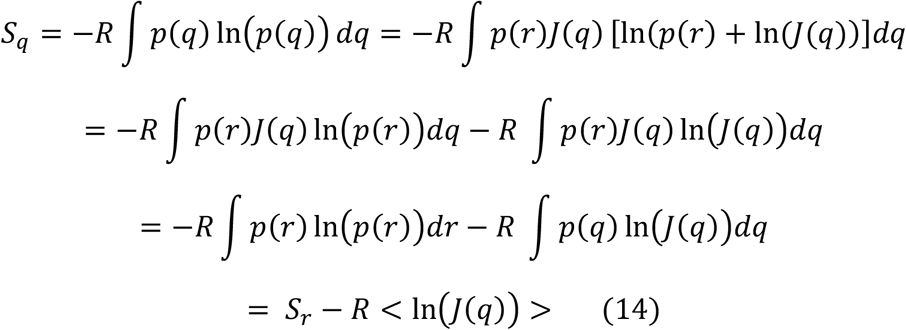

We then obtained the relationship of coordinate transformation:

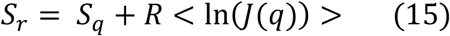

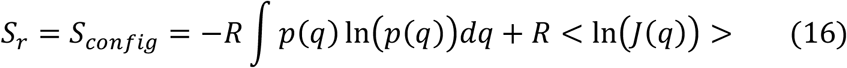

Transforming from Cartesian coordinate *r* ≡ (*x, y, z*) to BAT coordinate *q* ≡ (*b, θ, ϕ*), we can explicitly compute configuration entropy with the Jacobians 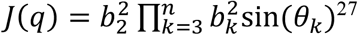

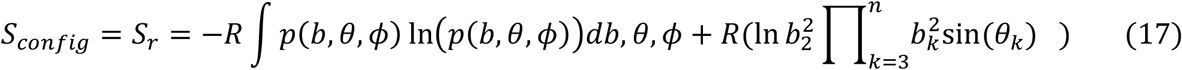

where *b*_*k*_ and *θ*_*k*_ are the bond length and bond angle of atom *k*, respectively.

#### Describing molecular configuration using PC modes with BAT coordinates

When a molecule has more than 4 atoms, directly using PDF in Gibbs’ entropy formula to compute S_config_ is not feasible. Internal BAT coordinate principal component analysis (iPCA) can be utilized for dimensionality reduction. Notably, we applied the same angle-aware arithmetic operation to avoid the discontinuity near ±π in all principal component (PC) calculations.

Standard mathematical protocols are employed to compute eigenvalues (*λ*) and eigenvectors (*v*) from iPC matrix. Notably, iPCA will result in 3N-6 eigen modes, where N is the number of atoms. All conformations sampled from MD simulations were projected onto the eigenvector of each eigenmode. This iPCA projection transforms the high-dimensional conformational space into a set of orthogonal one-dimensional distributions of an eigen mode. As an example, the integration of Gibbs entropy formula for each iPC mode can be written as *S*_*i*_:

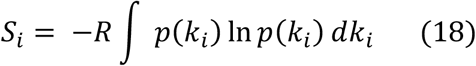

*k*_*i*_ and *p*(*k*_*i*_) represent the iPC coordinates and its corresponding PDF at eigenmode *i*, respectively.

The probability distribution of each PC is orthogonal to each other and thus the projected conformations of different eigenmodes are uncorrelated with each other. Therefore, the full PDF can be written as,

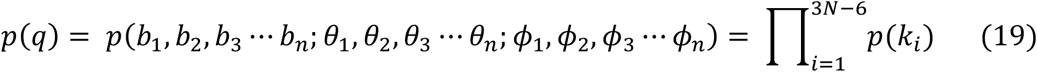

The total configurational entropy can be written as the sum of *S*_*i*_ with Jacobians

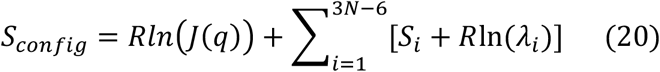

Eigenvalues (*λ*_*i*_) of eigenmode i represents the scaling (i.e. stretching or shrinking) along the eigenvectors, which is essential to correctly compute absolute molar entropies.

The integration of PDF (∫ *p*(*k*_*i*_) ln *p*(*k*_*i*_) *dk*_*i*_) can be evaluated numerically using basic quadrature. We can splice the PDF using histogram and compute the probability of each bin.

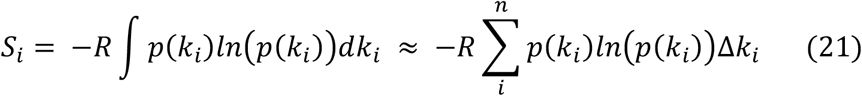

n represents number of bins. *p*(*k*_*i*_) represent the probability of each bin at each eigen mode. Δ*k*_*i*_ is the bin width of the histogram.

Know that integral of PDF = 1, ∫ *p*(*k*_*i*_) *dk*_*i*_ ≈ ∑ *p*(*k*_*i*_)Δ*k*_*i*_ = 1. Now consider the probability *p*′(*k*_*i*_) in each bin of the histogram, we know that

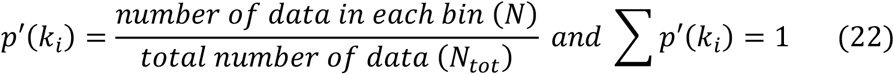

Knowing that ∑ *p*(*k*_*i*_)Δ*k*_*i*_ = ∑ *p*′(*k*_*i*_) = 1, we can easily derive the relation between *p*(*k*_*i*_) and *p*′(*k*_*i*_),

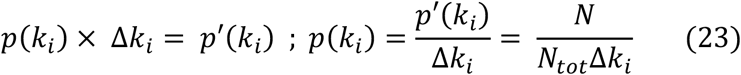

We can plug **equation (23)** back to **equation (21)**.

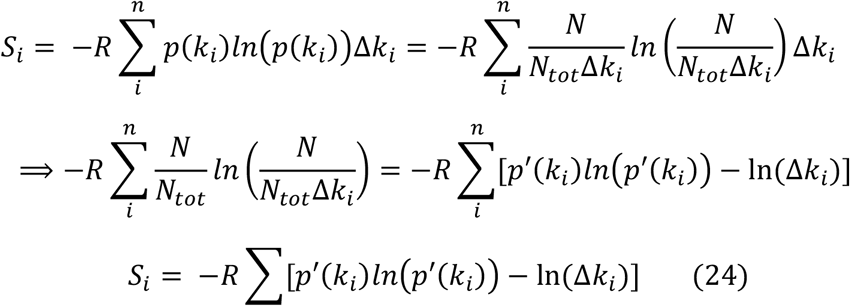

Plug **equation (24)** into **equation (20)** which shows the summation of configurational entropies (*S*_*i*_) at each eigenmodes, the Jacobian determinant *Rln*(*J*(*q*)) and the eigenvalues (*R*ln(*λ*_*i*_)) yield the full-dimensional configurational entropies (*S*_*config*_).

## Method

### MD simulations and molecular systems

All 3D conformations of alkanes and other organic molecules are obtained from the NIST website ^28^. MD simulations were performed using the AMBER22 package ^29^. The charges of the molecules were calculated using AM1-BCC ^30^and the force field parameters were assigned using the Generalized Amber Force Field (GAFF2) ^31^. All systems were solved in 15Å water boxes using TIP3P explicit solvent model at temperature of 298K with NPT ensemble ^32^. 12 Å cutoff was used for short range non-bonded interactions and the long-range electrostatic interactions were computed by the particle mesh Ewald method (PME) ^33^. The water molecules were minimized for 1000 steps, followed by minimization of the entire system for 2000 steps. The solvated system was equilibrated under constant pressure and temperature (NPT ensemble) from 50 K to 275 K with 25 K increments and 100-ps each, and finally at 298 K for 500-ps. Production runs were also performed in the NPT ensemble at 298 K using a Langevin Thermostat with 2-fs time-steps ^34^. We perform 200-ns MD simulation for all small molecules, and each frame was saved with 10-fs interval which made up a total of 20,000,000 frames. We resaved the trajectories every 1-ps which result in 200,000 frames for further analysis.

The initial structure of HIVp was taken from PDB 1HVR ^36^ where the co-crystalized Inhibitor XK2 and water molecules were removed. The protein was briefly minimized using MOE ^37^. In accordance with previous literature, HIS69/HIS168 were corrected to HIE and CSO 67/CSO166 changed to CYS ^38^. ASP25 was protonated to ASH, while ASP124 was not changed ^39^. RIT position was taken from PDB 1HXW ^40^. AMP position was taken from PDB 1HPV ^41^. Crystal structures 1HXW and 1HPV were aligned to 1HVR via pocket alignment using MOE. Inhibitor coordinates for AMP and RIT were then duplicated directly onto 1HPV. This results in identical protein coordinates for the apo, AMP bound, and RIT bound form, allowing us to compare entropy without adding additional bias. MD simulations were performed using the AMBER22 package ^30^ The charges of the molecules were calculated using AM1-BCC ^31^. 5 Cl-were added to neutralize the system. MD simulations used the ff14SB force field for protein ^42^ and GAFF2 for ligand ^32^. All systems were solved in 12Å water boxes using TIP3P explicit solvent model at temperature of 298K with NPT ensemble ^33^. 12 Å cutoff was used for short range non-bonded interactions and the long-range electrostatic interactions were computed by the particle mesh Ewald method (PME) ^34^. The water molecules were minimized for 10,000 steps, followed by minimization of the entire system for 20,000 steps. The solvated system was equilibrated under constant pressure and temperature (NPT ensemble) at 50 K for 200-ps, from 75 to 275 K with 25 K increments and 100 ps each, and finally at 298 K for 400-ps. Production runs were also performed in the NPT ensemble at 298 K using a Langevin Thermostat with 2-fs time-steps ^35^. We perform 500-ns MD saving every 1-ps interval. We then resaved the trajectories every 10ps which result in 50,000 frames. The first 7.5ns (750 frames) are treated as extra equilibrium and removed which result in a total of 49,250 frames for all following analysis.

### Translational and Rotational Entropy Calculations

The Translational and rotational entropies are solved analytically by integrating the 6 external degrees of freedom shown in **equations 3 and 4**. As shown in **equation 3**, translation entropy is determined by the volume of the molecules moved and thus it is subjected to the density of different molecules under a liquid environment. For instance, the density of water under standard condition is 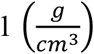, and the density is inversely proportional to the volume. The relationship between volume of water and volume of organic molecules can be calculated:

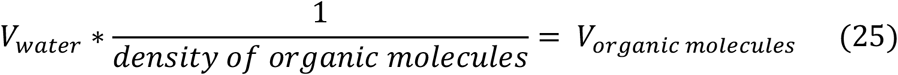

Therefore, the translational entropies of organic molecules become,

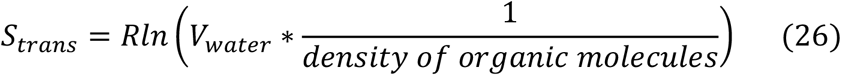

On the other hand, rotational entropies are determined by 8*π*^2^ and thus remain the same under liquid and aqueous environment as shown in **equation 4**. Then, the external entropy (*S*_*ext*_) liquid phase organic molecules can be written as:

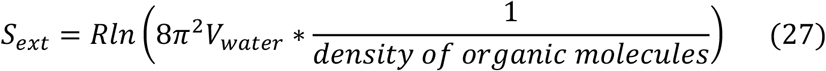

Known that the molar entropy (TS) of water is 4.98 (kcal/mol) ^35^, and the external entropies of organic molecules can be approximated as:

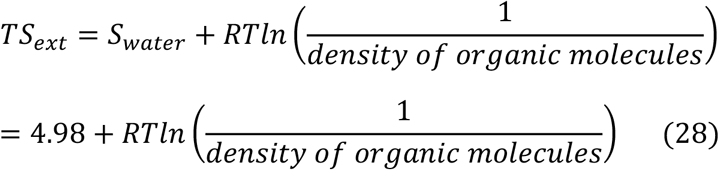

All density values of organic molecules are obtained from NIST Chemistry Webbook and PubChem ^28,36^.

In the case of HIV protease in complex with RIT and AMP, the systems are solvated in aqueous environment, standard molar concentrations 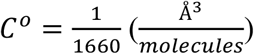 is used to account for the translational degree of freedom which is volume (*V*) in the solution under standard concentration ^37^.

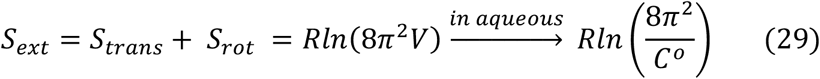

### Configurational Entropy Calculations

First, we extract the BAT coordinates of the molecules, then compute their eigenvectors and eigenvalues using iPCA with angle-aware arithmetic by mapping the torsion angles to (sin, cos) space when performing any mathematical operation. Next, standard statistical method was performed to obtain eigenvectors and eigenvalues from covariance matrix and then all input internal coordinates (200,000 conformations from 200-ns MD simulation for all organic molecules and 49,250 conformations from 500-ns MD simulation for HIVp protein-ligand binding) were projected on to the eigenvectors of each eigenmodes (**Figure S2 and S3**).

Projections were performed by computing the circular dot product between the BAT coordinate and their corresponding eigenvectors on all eigenmodes by using angle-aware mean and addition (**equation 10 and 11**). For the projection data on all eigenmodes, we approximate the probability distribution of the projected data using histogramming method. Once obtaining the probability of each bin under each eigenmodes, we can plug the probability (*p*′(*k*_*i*_)) and eigenvectors into the **equation (24)** which results in entropies of each eigenmodes.

In theory, the histogram method described in **equation (24)** estimates the probability of each microstate by dividing the coordinate space into bins. For PDFs that cover a large range of microstates, such as torsional rotations, the Gibbs entropy formula can correctly estimate the entropies. However, using Gibbs entropy formula can lead to unrealistic negative TS_config_ when a PDF appears to be the Dirac delta function (**Figure S4**). The delta distribution indicates a single and rigid state without molecular flexibility which does not contribute to TS_config_ (i.e. S ~ ln (1) = 0). As a result, for the iPC modes which present the uncorrelated single rigid state, we replaced negative entropy with 0 to avoid mathematics that leads to unphysical values.

Furthermore, the eigenvalues provide the scaling of each eigenmode, with smaller eigenvalues indicating smaller contributions. However, when only a small number of eigenvalues are present, the term ln(*λ*_*i*_) in **equation (20)** can become a large negative number. To avoid these artifacts, we discard all negative entropy contributions when summing over eigenmodes in **equation (20)**.

### Convergence Analysis

To obtain accurate results of configurational entropies calculation, we perform rigorous convergence analysis for all molecules. For organic molecules, we compute covariance matrix, eigenvectors, eigenvalues, projection and numerical for 1-2k, 1-4k, 1-6k … and 1-200k frames which result in 100 different values of configurational entropies for different number of frames used. Then we performed a convergence analysis to show the convergence behavior of TS_config_ **(Figure S5)**. In addition, we also show that the eigenvectors of pentane of the first 5 iPC remain stable after 160ns indicating it’s not necessary to recompute eigenvectors and eigenvalues when using longer simulation which can speed up calculation. (**Figure S6**).

#### Change in Entropy of Protein-Ligand binding: HIV-1 protease

We compute the change in molar entropy (*ΔS*) of the protein ligand complex: HIVp in complex with either RIT or AMP. We then compute the entropy of HIVp in complex with ligand (*S*_*R*+*L*_), entropy of free HIVp (*S*_*R*_), and entropy of free ligand (*S*_*L*_) separately.

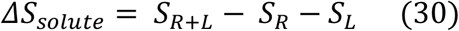

We know that the external and configurational terms can be computed separately.

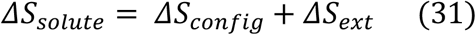

The change in configurational entropy (*ΔS*_*config*_) can be written into:

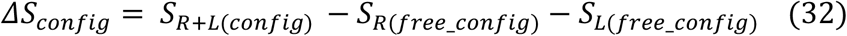

The configurational entropy of the bound form (*S*_*R*+*L*(*config*)_), can be further separated into the configurational entropy of protein bound (*S*_*R*(*bound*_*config*)_) and ligand bound (*S*_*L*(*bound*_*config*)_) by saving the MD trajectories with protein or ligand only.

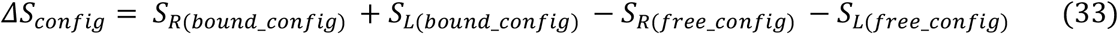

To enable comparison with published TΔS results with the VM2 method ^23^ and facilitate the calculations of *S*_*R*(*bound*_*config*)_ and *S*_*R*(*free*_*config*)_, we consider only residues within 7Å of the binding site. Residues outside of 7Å radius are not considered in the calculation (**Figure S1B**).

The change in external term (*ΔS*_*ext*_) is computed as:

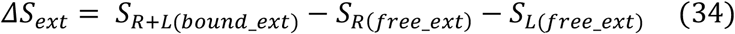

Similarly, *S*_*R*+*L*(*bound*_*ext*)_ can be broken down into external entropy of protein bound form (*S*_*R*(*bound*_*ext*)_) and ligand bound form (*S*_*L*(*bound*_*ext*)_).

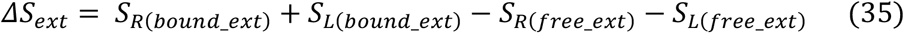

The external entropies of the protein in its bound (*S*_*R*(*bound*_*ext*)_) and free form (*S*_*R*(*free*_*ext*)_) are nearly identical and the difference can be approximated to zero. This leaves *ΔS*_*ext*_ from ligand bound and free form:

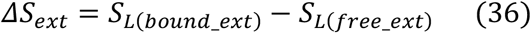

In aqueous environment, the external entropy term of the free ligand *S*_*L*(*free*_*ext*)_ is approximated using **equation (29)** under standard molar condition 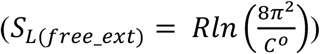. In ligand bound form, the bound ligand still can tumble and rotate within the binding pocket. Therefore, *S*_*L*(*bound*_*ext*)_ cannot be approximated as zero and our BAT internal coordinate system uses pseudo-bond that connected an atom in the ligand back to the root atom of protein to correctly account for *S*_*L*(*bound*_*ext*)_. As illustrated in **Figure S1A**, the pseudo-bond creates 6 extra degrees of freedom (1 bond, 2 angles and 3 torsions), and the translation and rotational motion along the pseudo bond can describe the tumbling motion of the ligand in the pocket. We first compute the covariance matrix, eigenvectors and eigenvalues of the 6 extra degrees of freedom, then we projected 6 extra degree of freedom along the eigenvectors for each iPC and computed the *S*_*L*(*bound*_*ext*)_ where 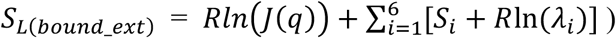. Alternative approaches by integrating all degrees of freedom (*S*_*ext*_ = *R* ∭ sin *θ dxdydzdrdθdφ*) will result in similar values (data not shown).

## Results and Discussion

### Molar entropy calculations for small organic molecules

This section begins by validating our computed molar entropy against available experimental data for small organic compounds ^38–44^. Notably, to enable direct comparison with energy, we include temperature 298K in our reported entropy calculations. **Table 1** and **Figure 3** compare the computed configuration entropy (TS_config_) and molar entropy (TS_comp_) with experimental molar entropy (TS_exp, 298K_) of a series of n-alkanes and other small organic compounds. Both TS_config_ and TS_comp_ accurately rank TS_exp, 298K_, thus validating the theory and computation strategies used in the iPC-entropy method as well as confirming that classical molecular mechanics force fields can accurately describe molecular motions. Our results show that TS_config_ is the major contributor to molar entropy and molecular flexibility. By examining each eigenmode, we can identify the most substantial torsional motions. Importantly, each PC mode must accurately represent the underlying molecular motion in order to yield reliable entropy estimates. When Cartesian coordinates are used, projecting the x, y, and z displacements of a torsional rotation produces linear atomic motions rather than a true angular rotation (**Figure 2B**), thus leading to inaccurate entropy calculations. Of note, torsional rotations have discontinuous boundaries at ±π; therefore, numerical operations must treat torsion angles appropriately to ensure correct arithmetic, as detailed in Methods.

**Table 1.** Experimental molar entropy (TS_exp,298K_) and computed molar entropy (TS_comp_) (kcal/mol)

| | $TS_{\text{config}}$ | $TS_{\text{ext}}$ | $TS_{\text{comp}}$ | $TS_{\text{exp},298\text{K}}$ | $TS_{\text{angle-restrained}}$ |
| --- | --- | --- | --- | --- | --- |
| Pentane | 9.76 | 5.45 | 15.21 | 18.73 | 13.89 |
| Hexane | 12.74 | 5.40 | 18.14 | 21.08 | 16.19 |
| Heptane | 14.13 | 5.36 | 19.50 | 23.36 | 18.37 |
| Octane | 17.29 | 5.33 | 22.63 | 25.71 | 20.57 |
| Nonane | 20.59 | 5.31 | 25.90 | 27.99 | 22.50 |
| 2-MethylHexane | 15.73 | 5.36 | 21.09 | 23.00 | N/A |
| 3-MethylHexane | 15.63 | 5.35 | 20.99 | 22.01 |  |
| Butanol | 9.16 | 5.19 | 14.35 | 16.08 |  |
| 1,2-Dichloroethane | 6.03 | 4.75 | 10.78 | 14.85 |  |
| Tert-Butyl alcohol | 9.38 | 5.20 | 14.58 | 13.50 |  |

$TS_{\text{config}}$ is solute entropy with the Jacobians included. $TS_{\text{angle-restrained}}$ was computed using MD trajectories with a restrained bond angle. External component $TS_{\text{ext}}$ was approximated using density at 298K. Notably, for easier comparison, our reported entropy includes temperature ( $T$ ) = 298 K, which yields the same unit (kcal/mol) as other thermodynamics data.

**Figure 3.**
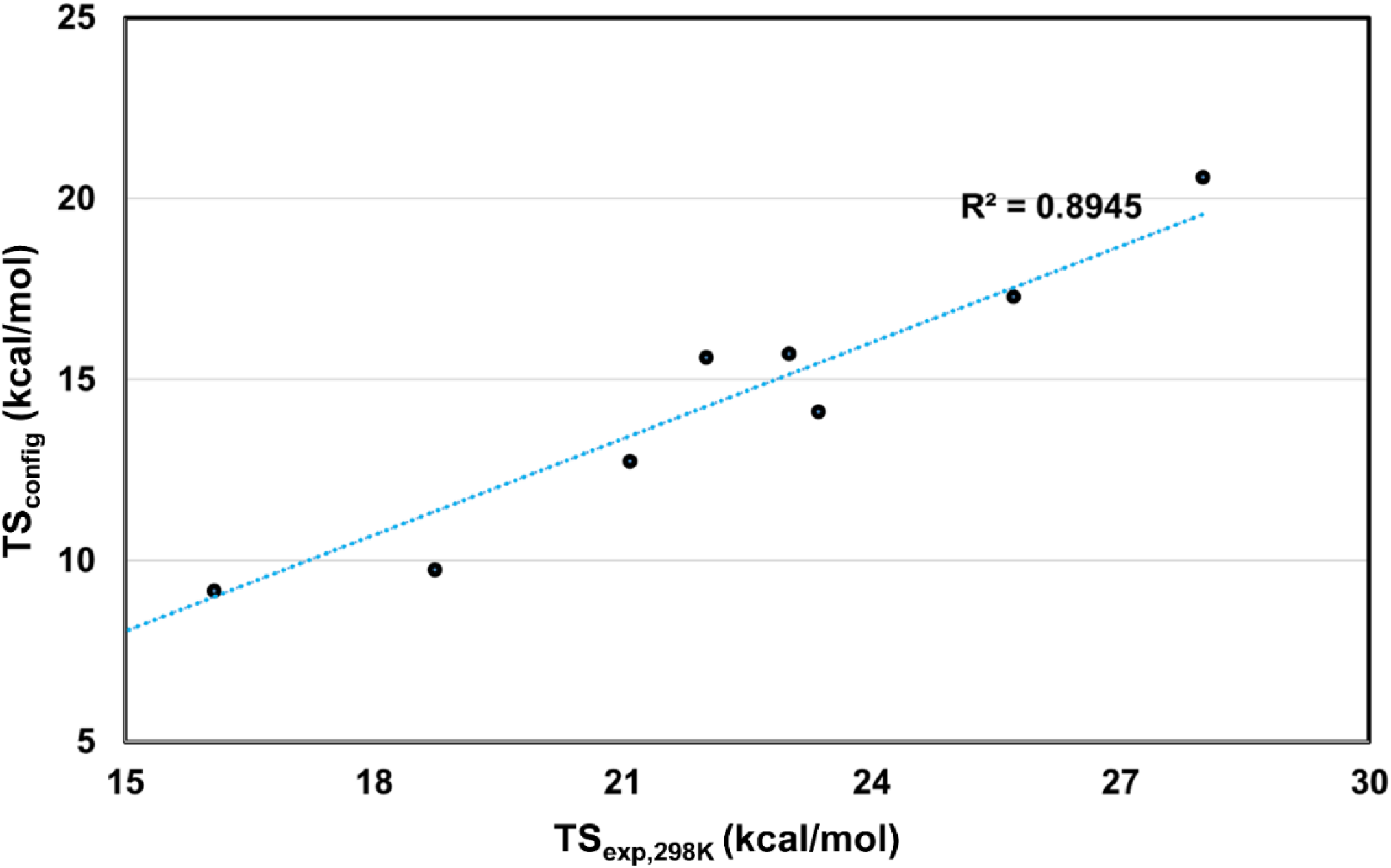
Evaluating the configurational entropy of the organic molecules. Comparing experimental molar entropy (TS_exp,298K_) and computed configurational entropy (TS_config_) values in kcal/mol. Linear fit and the corresponding correlation coefficient (R^2^ = 0.8945).

As illustrated in **Figure 3**, computed configuration entropy is highly correlated with molar entropy, with a nearly R = 1 correlation coefficient (R = 0.89). Note that TS_exp, 298K_ includes both a configuration component, TS_config_, from molecular flexibility, including molecular vibration and changing conformations, and an external component, TS_ext_, from their translational and rotational motions. This nearly perfect correlation indicates that internal motions are the main determinants of molar entropy. For all molecules, we found that TS_config_ was the main contributor to molar entropy, with more than half of molar entropy values from molecular internal motions (**Table 1**). Here we analytically computed TS_ext_ using **equation 27**, where the commonly used partition function considers all the translation and rotational degrees of freedom (DOF), which provide a reasonable approximation. Our results demonstrate that prioritizing accurate TS_config_ calculations is paramount. TS_comp_ values are consistently smaller than those of TS_exp, 298K_ (**Table 1**). The error may come from force field parameters in GAFF2, which may bring a systematic error, or from neglecting quantum effects. Nevertheless, we showed that using classical MD with the GAFF2 force field can correctly model small-molecule motions.

iPC-entropy approaches decoupled the BAT features into orthogonal modes. Then by projecting the data onto orthogonal eigenmodes, we obtained uncorrelated probability distribution across all eigenmodes. The projection of each eigenmode represents the linear combination of the BAT features, and its probability distribution describes substantial motion at each mode. We reduced a total of 3N-6 internal DOF to allow use of numerical integration of PDF to directly compute TS_config_ from each iPC mode (**equation 20**). Because PCA decouples molecular motions into independent orthogonal modes, we can sum TS_config,i_ computed from each iPC mode *i* to compute TS_config_ (i.e., **equation 24**). Notably, using the Gibbs entropy formula can lead to unrealistic negative TS_config_ when a PDF appears to be the Dirac delta function (**Figure S4**). The delta distribution indicates a single and rigid state without molecular flexibility, which does not contribute to TS_config_ (i.e., S ~ ln (1) = 0). As a result, for the iPC modes with the uncorrelated single rigid state, we replaced negative entropy with 0 to avoid mathematics that led to non-physical values.

The systematic increase in molar entropy observed in n-alkanes with increasing carbon number has been recognized and appreciated since the 1930s. For example, Finke and co-workers reported that the experimental molar entropy of n-alkanes C_8_ to C_16_ fitted a linear equation in N, the number of carbon atoms, where S_298.16K_ = 24.539 + 7.725N (cal/K·mol) or TS_298.16K_ = 7.316 + 2.303N (kcal/mol) ^41^. Our TS_comp_ of n-alkanes C_5_ to C_9_ yielded the same linear equation in N with a different offset, TS_comp_ = 2.175 + 2.59N. Both TS_298.16K_ and TS_comp_ of n-alkanes C_5_ to C_9_ fitted a linear equation in number of rotatable single bonds from carbon atoms (i.e., 2 to 6 for alkanes C_5_ to C_9_) (**Figure S7)**. Notably, except for linear alkanes, other small organic molecules are not likely to exactly fit the equation.

#### Key features contributing to TS_config_

In MD simulation analysis, the collective motions revealed by the first few iPC modes are often considered essential dynamics, referring to the most significant and/or functionally relevant movements. These motions are primarily driven by torsion angle rotations, as illustrated in **Figure 4**. Mathematically, these iPC modes are ranked according to the variance captured by their corresponding eigenvectors, which also quantify the coverage of the motions among the overall conformational fluctuations. Therefore, we anticipated that the computed TS_config_ values would be linearly correlated with their eigenvalues (i.e., the first mode yields the largest computed TS_config_). In contrast, the computed TS_config_ from each iPC mode was not linearly proportional to its eigenvalue (**Table S1**), and iPC modes with significant torsion rotations near the center of a molecule typically resulted in large TS_config_, although they are not the first couple of iPC modes. For example, in hexane, the first 3 iPC modes contributed equally to TS_config_, and the third iPC mode of octane had the largest TS_config_ (**Table S1**). Estimating which iPC mode contributes most to the configuration entropy without performing numerical integration (**equation 24**) and properly accounting for the scaling factor is not straightforward. As demonstrated in nonane, only the first iPC mode showed distinct rotational states in the histogram plot (**Figure 4**), but the second iPC mode yielded the largest TS_config_. A similar pattern was observed for the drugs RIT and AMP, where the first 6 iPC modes each contributed ~ 2 kcal/mol of TS_config_, regardless of their corresponding eigenvalues (**Table S2**). These results highlight the significant contribution of the central torsions to molecular flexibility, which is reflected in the computed TS_config_. As shown in **Figure 4 and S8**, PDF is presented as histograms for configuration distribution based on eigenvectors. Distorting conformation along eigenvectors provides further information for the concerted motions and their contribution to TS_config_. The first iPC mode clearly depicts methyl rotations in the edges of the molecules. However, torsion angles centered on a molecule from the third iPC mode and their correlation with other DOF (i.e., angles) are the key determinants of molecular motions (**Table S1**), which can be quantified with TS_config_ using **equation 20**.

**Figure 4.**
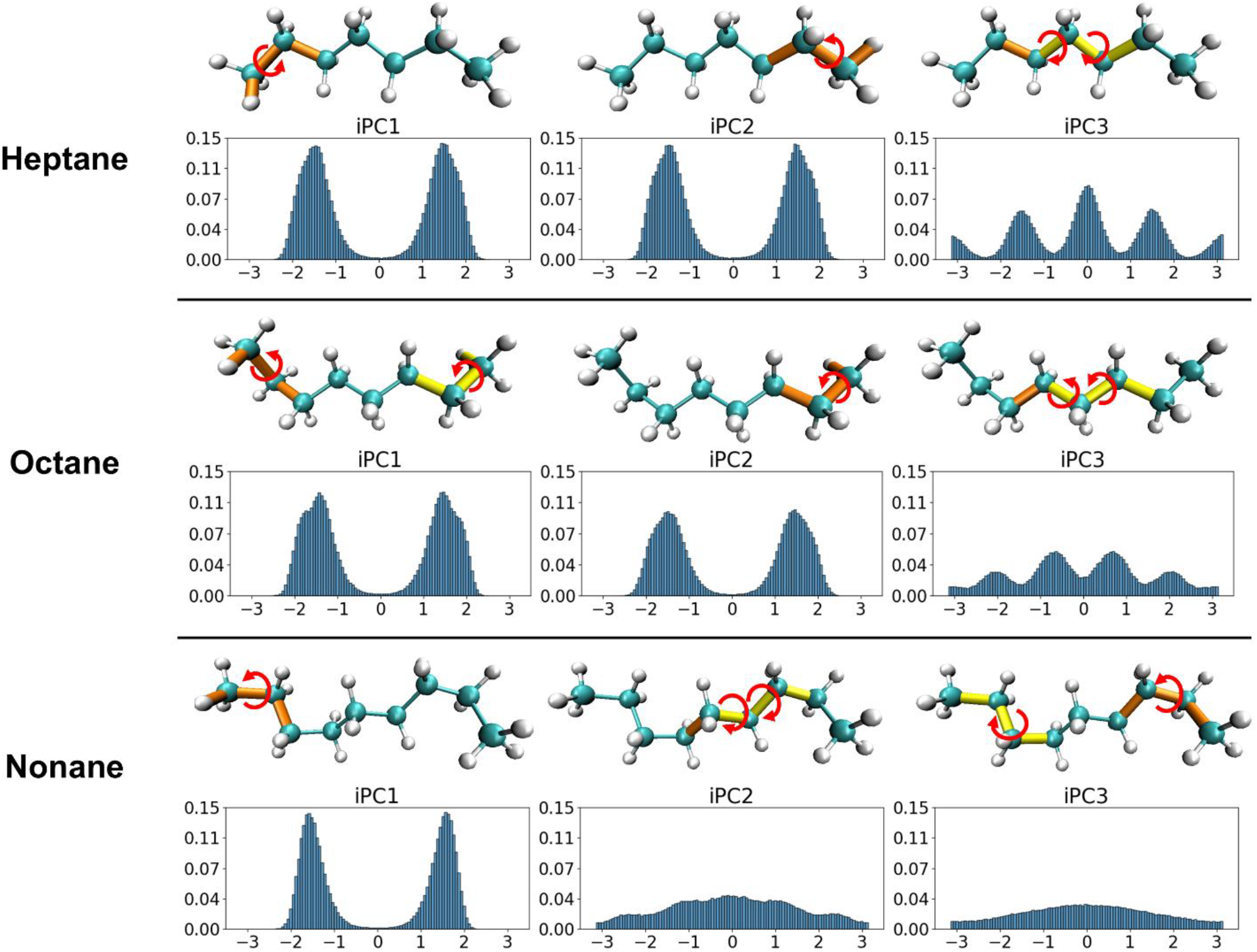
Dissecting molecular motion of n-alkanes C_7_ to C_9_ of the first three internal-coordinate PC-entropy (iPC) modes. The probability distribution of projection at each iPC mode is shown, with major torsion rotation highlighted. Red and yellow indicate the top two torsion rotations and the red circular arrow represents the direction of rotation.

#### Rotamer-counting approaches. Can rotatable bonds be used without considering angle bending to accurately compute TS_config_?

Single (σ-) bonds connect atoms and allow atoms to rotate; they are commonly called rotatable bonds or rotamers. Here we call it primary torsion (PrimaryTor). Because analyzing PrimaryTor is straightforward, we post-processed MD trajectories by considering only these rotatable bonds to construct individual PDFs of each PrimaryTor to compute the Gibbs entropy (TS_PrimaryTor_). Surprisingly, summing rotatable bond entropy TS_PrimaryTor_ calculated by each individual rotamer without accounting for their correlations yielded a value smaller than that of TS_config_ (**Table S3**). This finding contradicts intuition because neglecting correlations is expected to produce an upper bound on entropy. However, because linear alkanes do not have pronounced intramolecular attractions, correlations between the torsions may be minimal. In addition, we found that TS_PrimaryTor_ = 0.009 + 1.379N (kcal/mol), with the linear relationship different from TS_298.16K_ and TS_comp_, thereby indicating that traditional entropy calculations focusing on merely rotatable bonds may overlook important contributions from angle bending. The full configurational space corresponds to the volume accessible to the molecule in the internal coordinate space. Because the configurational integral formally includes integration over both dihedral and angle DOF, excluding angle bending would substantially reduce the accessible configurational space.

Angles modulate the spatial range available for dihedral rotations and therefore cannot be neglected. To examine how angles contribute to molecular flexibility and TS_config_, we carried out another set of MD runs with angular constraints. Using pentane as an example in **Figure 5A and S9**, all torsion angles sampled the full rotational range (i.e., from −π to π), although their population distributions varied slightly. Although angle bending did not alter the torsional ranges, it could couple with torsional motion to enable the molecule to fully explore configurational space (**Figure 5B**). Restraining angle bending diminished angle-torsion coupling, thus resulting in decreased flexibility and losing 1 to 3 kcal/mol to TS_config_ **(Table 1)**. However, by using the same histogramming method, selecting only angle-bending DOF in eigenvectors resulted in a sharp peak/single bin (**Figure S10)**, so angle bending alone cannot contribute to conformational changes of n-alkanes. Of note, the strength of angle–torsion coupling grew non-linearly with increasing carbon chain length in n-alkanes (**Table 1**). These calculations highlight the critical role of bond angles in coordinating with torsional rotations, enabling molecules to access the full spectrum of configurational space.

**Figure 5.**
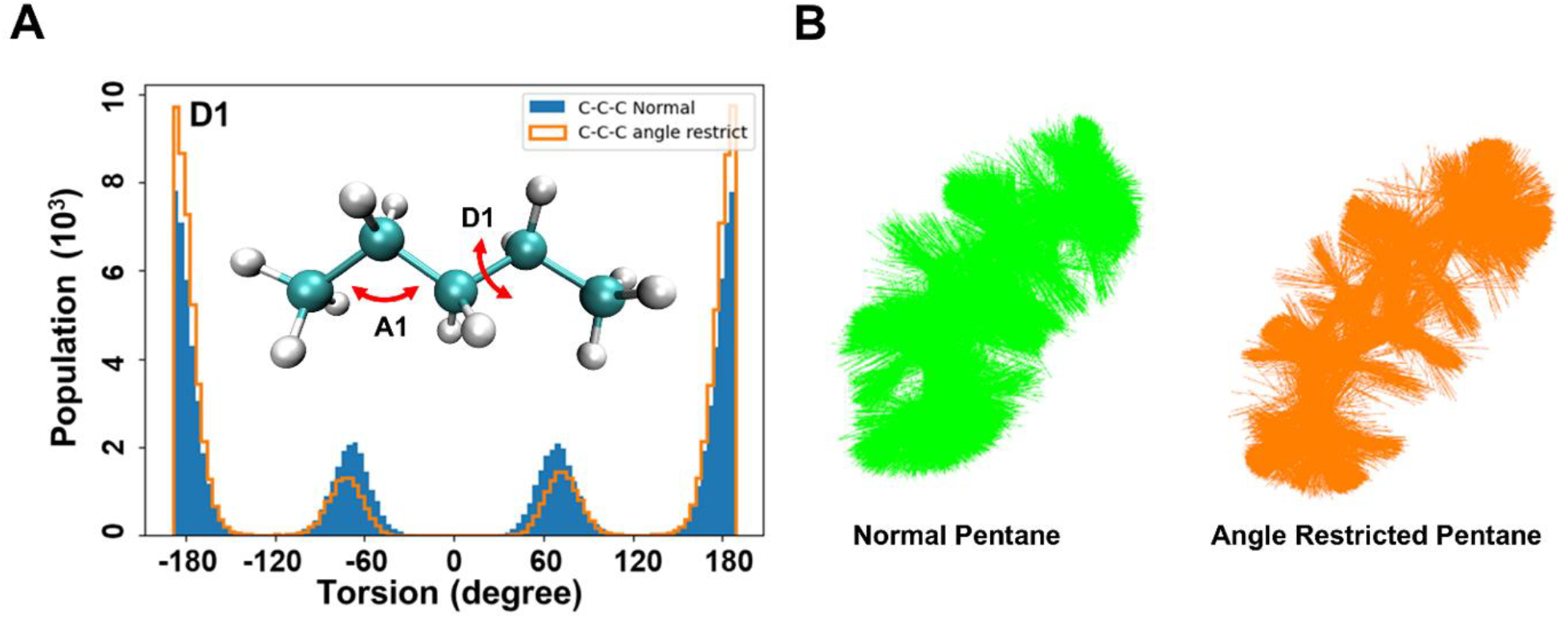
Observing the angle contribution to entropy. **(A)** Probability distribution of dihedral angle, D1, of normal pentane and angle-restricted pentane. **(B)** Overlaying the 1000 frames of molecular dynamics trajectories of normal pentane and angle-restricted pentane.

### Loss of solute entropy upon drugs binding to HIVp

Post-processing classical MD trajectories, free HIVp, free drug, and drug-HIVp bound complex, with focusing on a drug and residues of proteins within 7 Å of the bound drug with iPC-entropy, the computed TΔS°_Solute_ showed that binding RIT and AMP to HIVp lost 37.72 and 30.84 (kcal/mole) solute entropy, respectively (**Table 2**). The computed entropy losses are in good agreement with results computed with the VM2 program ^23^, a rigorous method to calculate free-energy G° in standard concentration. The configuration entropy is computed by TS°_Solute_ = <U+W> - G°, where U is the molecular mechanics energy and W is the solvation free energy. Because W includes both the solvent enthalpy and solvent entropy terms, the equation yields configuration entropy TS°_Solute_, which cannot be directly compared with experimentally measured entropy changes. To accelerate G° calculations, VM2 samples conformations of the bound ligand and protein residues within 7 Å of the bound ligand. For a fair comparison, we selected 42 HIVp residues within 7 Å of bound RIT or AMP to compute S°_Solute_ using the iPC-entropy method. Note that both methods use **equation 29** to compute external entropy S_ext_ for each free species, which is TS°_ext_ = 6.98 kcal/mol under standard conditions at 298K. The values of total entropy changed TΔS°_Solute_ include the decrease in the drug’s translational and rotational mobility in the bound form, and both drugs reducing TS_ext,bound drug_ to 1.83 kcal/mol (**Table 2**) (see Methods). Nevertheless, different ligands may lose rigidity upon binding in various degrees, and their TΔS_ext_ should not be assumed as a fixed value.

**Table 2.**
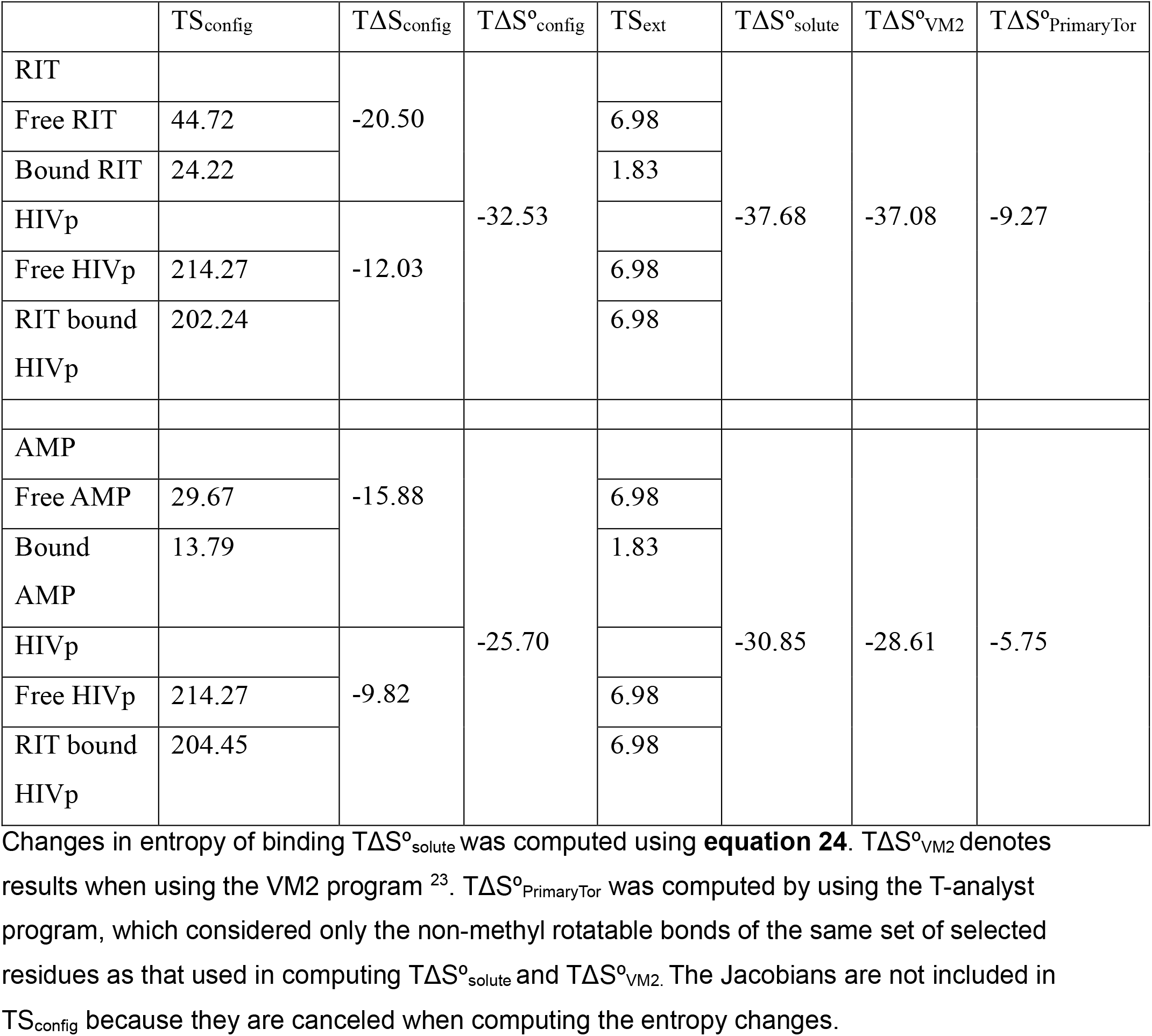
Computed solute entropy TS_solute_ of each species and changes in TΔS°_solute_ on ritonavir (RIT) and amprenavir (AMP) binding to HIV protease (HIVp) in standard concentration (kcal/mol).

The large TΔS°_Solute_ is not surprising because both drugs tightly bind HIVp. Notably, although experimentally measured binding free energy ΔG is ~ −13.5 kcal/mol for both drugs ^45,46^, RIT binding to HIVp yields ~7 kcal/mol more configuration entropy lost than AMP binding to HIVp. The results highlight the importance of computing TΔS°_Solute_ to accurately predict ligand binding affinity. Because TΔS_ext_ only accounts for ~5.15 kcal/mol entropy loss, the main source of TΔS°_Solute_ is from rigidifying molecular configuration upon binding, which strongly opposes binding RIT and AMP to HIVp, with TΔS_config_ contributing to −32.53 and −25.70 (kcal/mol), respectively (**Table 2**). Of note, the major contributor is from losing the configuration entropy of the drug rather than the protein. As shown in **Table 2**, TΔS°_Solute_, RIT (i.e., 98 atoms, 18 rotatable bonds) lost 20.50 kcal/mol configuration entropy, and AMP (i.e., 70 atoms, 12 rotatable bonds) lost 15.88 kcal/mol configuration entropy upon binding to HIVp. However, the binding pocket of HIVp has a total of 42 residues and 138 non-methyl rotatable bonds from the backbone and sidechains; these residues lost only 12.03 and 9.82 kcal/mol configuration entropy when binding with RIT and AMP, respectively. Sidechains of the HIVp binding pockets are restricted even in the free state because of the confined space. In contrast, free ligands possess much greater flexibility (**Figure 6**). Therefore, the entropic loss from rigidifying dihedral rotations upon binding is far more pronounced in the ligand.

**Figure 6.**
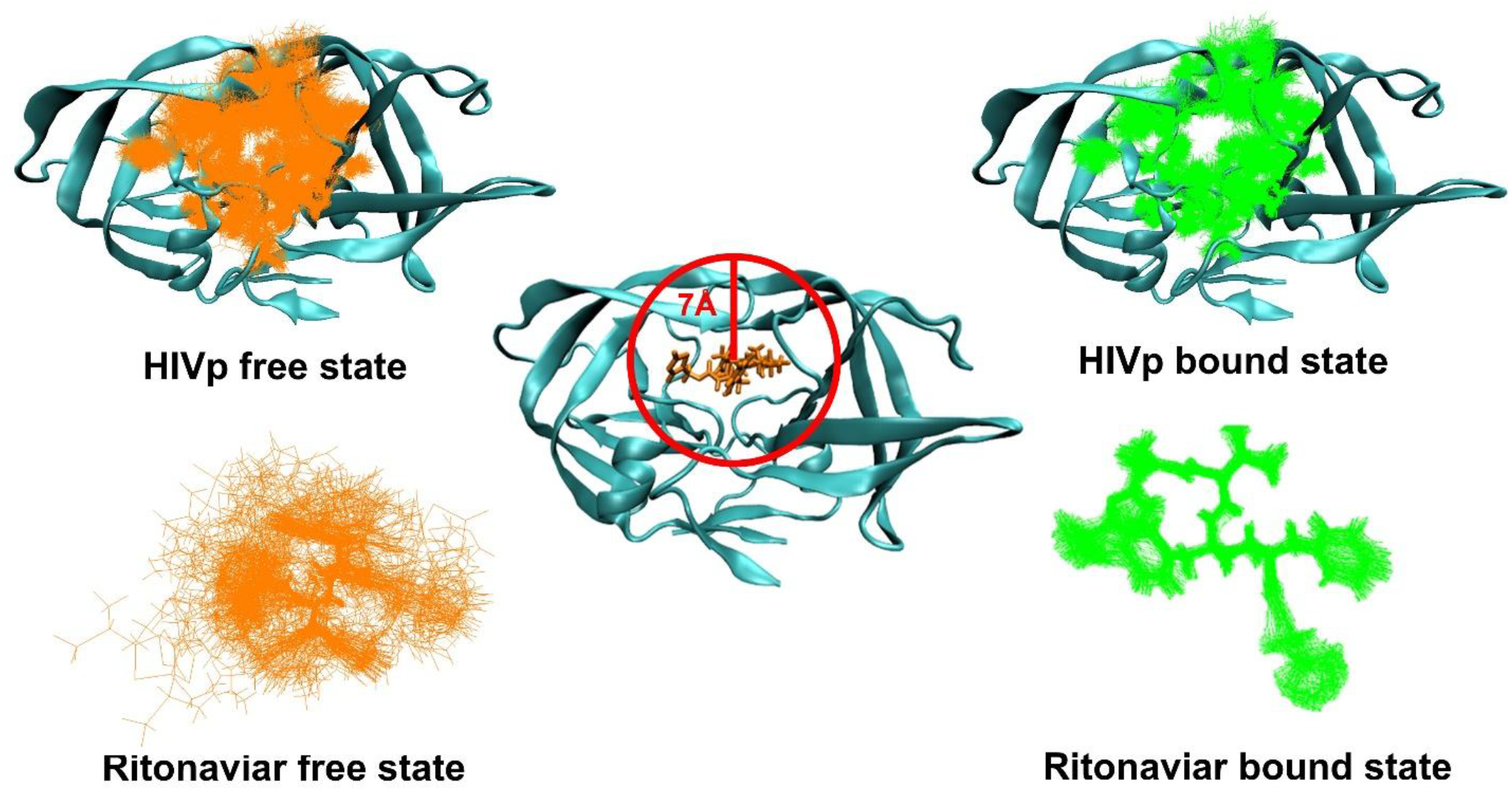
Changes in configurational dynamics of ritonavir in the complex of HIV protease (HIVp) and RIT in the free state. (**Center**) HIVp is shown in cyan with 7 Å life set selected from the center of ritonavir. (**Left**) Overlay of 100 frames of molecular dynamics (MD) simulation of RIT and HIVp free state. (**Right**) Overlay of 100 frames of MD simulation of RIT and HIVp bound state. HIVp is shown in cyan.

Figure 6. illustrates that upon binding to HIVp, each rotatable bond in RIT is restricted to a single rotamer state with remarkably narrower rotational ranges, especially for the torsions near the center of RIT. Torsion rotations can be strongly coupled with tiny angle bending (i.e., 1°-2°) to significantly expand the overall molecular flexibility. Therefore, we anticipated that restricting angle bending would rigidify the drug’s motion and reduce the entropy penalty upon binding. We performed another set of MD simulations with angle constraints on the drugs. Both drugs had smaller TS_angle-restrained_ in their free and bound forms, and RIT showed ~7 kcal/mol entropic loss upon binding to HIVp (i.e., TΔS_angle-restrained_ = −13.82 kcal/mol and TΔS= −20.50 kcal/mol) (**Table S4**). Of note, we found no reduction in entropy loss by rigidifying AMP. Rigidified ligands unexpectedly restrict HIVp mobility upon ligand binding, thus driving a larger entropy penalty due to the loss of protein sidechain flexibility. Although the pronounced entropy loss of HIVp during RIT binding was offset by the ligand’s minimal conformational entropy loss, reducing the net entropic penalty by 2 kcal/mol, this compensation was absent for AMP. Instead, the reduced flexibility of HIVp induced by rigidified AMP yielded 2 kcal/mol greater overall entropy penalty. Our study underscores that contrary to the common assumption that rigid drugs can effectively minimize binding entropy losses, they can significantly restrict protein dynamics, thereby counterproductively hindering binding.

### What is missed in entropy calculation from estimating torsion entropy using rotatable bonds?

A popular approach to approximating the entropy penalty of protein–ligand binding in molecular docking and scoring functions is to use a penalty proportional to the number of rotatable bonds (PrimaryTor). Using pentane as an example, the molecule has two rotatable bonds (rotamers) between C_2_-C_3_ and C_3_-C_4_. Atoms C_1_, H_1_ and H_2_ (i.e., C_1_-C_2_-C_3_-C_4_ and H_1or2_-C_2_-C_3_-C_4_) all share the same single bond between C_2_-C_3_, and we used 4 heavy-atom C_1_-C_2_-C_3_-C_4_ torsion to define this PrimaryTor. Rotating this bond can move atoms C_1_, H_1_ and H_2_ in concert. Post-analysis of MD trajectories using each PrimaryTor does not need iPCA to construct the PDF for Gibbs entropy calculations. However, although this post-analysis is simple, as illustrated in **Table 3**, TS_PrimaryTor_ largely underestimates configuration entropy. These errors do not cancel when computing the entropy loss upon binding, yielding TΔS_PrimaryTor_ values of −9.27 kcal/mol and −5.75 kcal/mol for RIT and AMP, respectively, and TΔΔS_PrimaryTor_ of ~ 4 kcal/mol, which affects the accuracy of binding free energy calculations. Notably, the TΔS°_solute_ was −37.68 kcal/mol for RIT as compared with −30.85 kcal/mol for AMP (**Table 2**), yielding TΔΔS °_solute_ of ~ 6.8 kcal/mol.

**Table 3.** Computed entropy TS_PrimaryTor_,TΔS°_PrimaryTor_, and TΔS°_PrimaryTor+correct_ on ritonavir or amprenavir binding to HIV protease (HIVp) in standard concentration (kcal/mol).

| | $TS_{\text{PrimaryTor}}$ | $T\Delta S_{\text{PrimaryTor}}$ | # of Flexib<br>Primary Torsion<sup>o</sup> | $S_{\text{correct}}$ | $TS_{\text{PrimaryTor+correct}}$ | $T\Delta S^{\circ}_{\text{PrimaryTor}}$ | $T\Delta S^{\circ}_{\text{PrimaryTor+correct}}$ |
| --- | --- | --- | --- | --- | --- | --- | --- |
| Free RIT | 7.79 | -5.59 | 17 | 5.78 | 13.57 | -9.27 | -17.43 |
| Bound RIT | 2.20 |  | 6 | 2.04 | 4.24 |  |  |
| Free HIVp | 11.84 | -3.68 | 51 | 17.34 | 29.18 |  |  |
| Rit bound HIVp | 8.16 |  | 38 | 12.92 | 21.08 |  |  |
| Free AMP | 6.31 | -4.34 | 13 | 4.42 | 10.73 | -5.75 | -8.13 |
| Bound AMP | 1.97 |  | 6 | 2.04 | 4.01 |  |  |
| Free HIVp | 11.84 | -1.41 | 51 | 17.34 | 29.18 |  |  |
| AMP bound HIVp | 10.43 |  | 51 | 17.34 | 27.77 |  |  |

$TS_{\text{PrimaryTor}}$ was computed using the T-analyst program, which considered only the non-methyl rotatable bonds. Primary torsion with $>30^{\circ}$ are counted as flexible primary torsion, where $S_{\text{correct}} = 0.34$ is applied only to those torsions. Primary torsion angles with $< 30^{\circ}$ of movement result in near-0 entropies and thus are not considered for correction. $T\Delta S^{\circ}_{\text{PrimaryTor}}$ and $T\Delta S^{\circ}_{\text{PrimaryTor+correct}}$ represent the overall entropy change of binding with and without $S_{\text{correct}}$ . External entropies are not included. The Jacobians are also not included because they are canceled when computing the changes.

We dissected the error source of TS_PrimaryTor_ and compared with TS_Solute_ to identify the systematic differences for correction. Using an MD run of pentane, rotatable bond C_2_-C_3_ defined by the heavy atoms C_1_-C_2_-C_3_-C_4_ yielded TS_PrimaryTor_ of ~ 0.67 kcal/mol. However, inspection of the MD trajectory revealed that the attached atoms (C1, H1, and H2) do not rotate perfectly in concert because of bond-angle fluctuations and other internal motions. For instance, when C_1_ moved 60°, H_1or2_ may move 55° or 65°. We computed the circular correlation coefficients (ρ) using angle-aware arithmetic (detail in Supplementary material) between torsions sharing the same rotatable bond. The circular correlations were typically 0.8–0.9 rather than the ideal value of 1. Because these torsions remained highly correlated, their joint distribution can be approximated as multivariate Gaussian. Under this approximation, the correlation between two torsions is quantified by their mutual information,

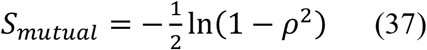

Mutual information (*S*_*mutual*_) represents the amount of entropy shared between correlated torsions and therefore corresponds to the entropy that would otherwise be counted multiple times if each torsion were treated independently. For example, when ρ=0.8, the mutual information is approximately 0.5 kcal/mol. Because a typical primary torsion contributes approximately 0.67 kcal/mol, the remaining independent entropy associated with an additional correlated torsion is only about 0.17 kcal/mol. A typical primary torsion is coupled to two secondary torsions (e.g. H_1or2_-C_2_-C_3_-C_4_), so neglecting these secondary torsions omits approximately 0.34 kcal/mol of entropy per primary torsion. Therefore, we used 0.34 kcal/mol as a correction factor (*S*_correct_) for each PrimaryTor, with flexible primary torsion defined as primary torsion with movement > 30°. Although this value may slightly overestimate the correction because some primary torsions are coupled to only one secondary torsion, this method served as a decent approximation. Applying *S*_correct_ to all PrimaryTor yields 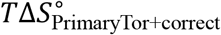 values of −17.43 and −8.13 kcal/mol for RIT and AMP binding, respectively (**Table 3**). Notably, this simple correction helps explain why the observed difference in solute entropy, 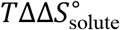, is ~ 8.5 kcal/mol, whereas the uncorrected primary torsion entropy difference, 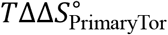, is only ~ 3.5 kcal/mol (**Tables 2 and 3**). After applying the correction, 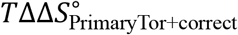 increases to approximately 9 kcal/mol, in good agreement with both the iPC entropy and VM2 estimates (**Table 2**). Unlike the PrimaryTor rotamer-counting approach, the iPC-entropy method explicitly accounts for correlated motions via its eigenvectors and eigenvalues, which capture dimensionality reduction in configurational space and provide a more complete description of configurational entropy. Consequently, relying solely on primary torsion entropy without accounting for correlated motions can lead to substantial entropy errors and potentially incorrect rankings of ligand binding affinities, which may adversely affect drug discovery efforts.

### Analysis of computation cost for iPC-entropy calculations

The overall configurational entropy calculation time was < 5 s and 10 min for small molecules with 10,000 and 100,000 frames, respectively, including n-alkanes and the drugs RIT and AMP. Using the same single Intel Xeon E3-1246 CPU core for HIVp, the calculation time was approximately 1.5 hr with 10,000 frames. The most computationally expensive step was projecting the MD frames onto the eigenvectors. The computational cost scales linearly with both the number of MD frames and the number of BAT features (equivalent to the number of iPC modes). For example, projecting 10,000 and 50,000 MD frames onto 285 iPC modes required approximately 1.5 hr and 7.5 hr, respectively, on this single CPU core. Numerical integration of the 1D PDF for each iPC mode is relatively inexpensive, taking 2 and 10 s to process 10,000 and 50,000 data points, respectively, for each iPC mode of HIVp using the same single CPU core.

We demonstrated that use of 10,000 MD frames is sufficient to yield a converged entropy value for drug-protein systems. As shown in **Figure S11**, the 1D PDF, a 50k-datapoint histogram obtained using an eigenvector from 50,000 frames, shows reduced bin-to-bin variance/statistical noise relative to the 10k-datapoint histogram because a larger sample gives a lower standard error per bin (approximately scaling as 1/√N). Nevertheless, the overall shape and width of each distribution are nearly identical between histograms generated using 10,000 and 50,000 MD frames, so the projections along each iPC using 10,000 frames had already converged, and the extra frames primarily improve the precision of the density estimate rather than changing its computed TS_config_ (**Table S5**). Of note, if the data points are insufficient to generate a smooth distribution for integration, one may use interpolation or kernel density approximation to smooth the probability distribution, as shown in Supplementary material. The adaptive quadrature methods can be applied to dynamically adjust the bin width.

## General comments and conclusions

This study describes iPC-entropy, a new computation approach to post-process MD trajectories for accurately computing molar entropy and molecular configuration entropy. Following the nature of molecular motions and energy equations used in MM force fields, we demonstrated that using eigenvectors from PCA with BAT coordinates can correctly describe the molecular motions and ensembles of molecular configurations for iPC-entropy to perform entropy calculations with the Gibbs entropy formula.

Our computed molar entropy (TS_comp_) can reproduce experimental molar entropy measurements (TS_exp,298K_) with approximately 0.9 correlation, so MM force field GAFF2 may accurately describe molecular motions for n-alkanes and other organic molecules. The calculations also reveal the key features contributing to configuration entropy, including unexpected observations about the critical role of angle bending to allow coordinated angle bending and dihedral rotations in configuration space. Also, angle-torsion coupling increased non-linearly with the size of the molecules.

Analyzing entropic penalty TΔS_config_ upon drugs binding to the target HIVp shows that rigidifying the ligand and residues within the binding pocket can contribute to >25 kcal/mol unfavorable binding, which largely compensates the intermolecular attraction enthalpy, thereby highlighting the importance of computing TΔS_config_ accurately to predict binding affinity. Substantially narrowing the configurational space of the center dihedral of the drugs mainly contributes loss in the drug’s entropy loss. Notably, our iPC-entropy protocol can also post-analyze MD trajectories obtained from enhanced sampling techniques or machine learning to estimate TS_config_. However, if the population of their conformational free energy landscape is perturbed, one may need to reweigh TS_config_ to ensure accurate TS_config_ and TΔS_config_ calculations.

In computer-aided drug discovery, estimating torsional entropy loss by counting rotamers (primary torsions) is a common approach used in molecular docking and scoring functions. Our results show that torsional entropy computed by post-analyzing only rotamers in MD trajectories (TS_PrimaryTor_) substantially underestimates entropy loss, which in turn can produce large errors in binding affinity predictions. We revealed the fluctuations ignored by rotamer-counting approaches and added a correction term to systematically correct TS_PrimaryTor_. Although the absolute entropy loss (TΔS_PrimaryTor_) is less accurate than using all BAT DOF (TΔS_Config_), the relative entropy changes after correction 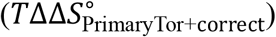 reliably reproduce the relative entropic penalty and thereby preserve accurate ranking of drug binding.

In summary, the present calculations indicate that the iPC-entropy method can accurately evaluate both the absolute molar entropy and entropic penalty upon protein–ligand binding from MD trajectories. Because MD simulations are a popular method for studying molecular motion, our approach is particularly valuable in accurately quantifying molecular flexibility via post-analysis of MD trajectories. iPC-entropy allows for more accurate protein–ligand binding affinity prediction and reveals key dihedral motions, which opens a new avenue of future drug and protein design.

## Data and software availability

The code and data sets are available on https://github.com/chang-group

## Acknowledgements

We thank Dr. Zhiye Tang for helpful discussion. This study was supported by the US National Institutes of Health (R01GM-109045 and R35GM-164210 to C. C.), the US National Science Foundation (MCB-2437134 to C.C.), and a UCR RAISE fellowship to T.H., UCR Academic Senate fellowship to C. C. and Department of Education fellowship for E.V.

## Notes

### Competing Interest Statement

The authors have declared no competing interest.

