## Supporting Information for "Mechanistic Dissection of Entropic Penalty upon Ligand Binding and Molecular Flexibility via Molecular Dynamics Simulations and Machine Learning"

**Smoothing the Probability Distribution with Kernal Desnity Approsimation**

Sufficient datapoints (i.e. larger than 1000 datapoints) are required while performing numerical integration to compute configuration entropy using Gibbs' entropy formula for each iPC mode. If a system has few datapoints which cannot generate continuous probability distribution in a histogram plot, smoothing the probability distribution is necessary to perform numerical integration. To achieve this, we can apply the weight based on gaussian kernel density estimation (KDE) to approximate the probability distribution of our data. Given a dataset with n data points, a kernel function K(x) and bandwidth (σ). The kernel density estimate (KDE) is defined as,

$$f(x) = \frac{1}{n\sigma}\sum_{i=1}^{n} K(\frac{x - x_i}{\sigma}) \qquad (eq\ S1)$$

$x_i$ represent the input data. $x$ is the grid of the data. n is the number of data points. σ represents the kernel bandwidth which is the distance between each kernel. The larger the bandwidth the smoother the resulting distribution; smaller bandwidth can capture more detail. Therefore, there is a trade off of how smooth you want your distribution while avoiding capture noise of the data. In this study, bandwidth **(σ = 0.05)** was used to produce smooth distribution while not losing information.

For a gaussian kernel,

$$K(x) = \frac{1}{\sqrt{2\pi}} e^{-\frac{x^2}{2}} \qquad (eqS2)$$

After obtaining probability distribution by KDE, we provide an inverse relationship with probability and bin width as written below:

$$bin\ width\ (w) = \frac{\left(1 - f(x)\right)}{\sum\left(1 - f(x)\right)} 2\pi \qquad (eqS3)$$

**Detailed examination of computation time and numerical integration**

The entropy calculation for small molecules n-alkane and drug-like compounds was computed in less than 10 minutes. For HIVp, this calculation took approximately 1.5 hours on a single Intel Xeon E3-1246 CPU core. The most time-consuming step is projecting MD frames onto each eigenvector. The computation time scale linearly with number of MD frames and the number of BAT features which correspond to number of PC components. For example, projecting 50,000 MD frames onto 285 eigenvectors, each eigenvector has 285 components, took ~7 hours with the single Intel CPU core. However, other post-processing analyses performed on the same single CPU core can be slower; for example, computing Molecular Mechanics Poisson-Boltzmann Surface Area (MM/PBSA) calculations for the same 50,000 frames took longer than 200 hours.

Numerical integration of each 1D probability distribution in each iPC mode is not time consuming. It took ~1hours and ~10 minutes for 50,000 frames and 10,000 frames, respectively. We examine if the smoothness of the probability distribution will affect the $TS_{config}$ of numerical integration by projecting the 10,000 MD frames onto the eigenvectors obtain from 50,000 or 10,000 frames. As illustrate in **Figure S11**, the probability distribution of 50,000 frames is smoother than 10,000 frames but the computed configurational entropy yield nearly identical $TS_{config}$ (**Table S5**).

If the data points are insufficient to generate a smooth distribution for integration, one may use interpolation or kernel density approximation to smooth the probability distribution, as exampled in Supplementary material. Adaptive numerical integration, also termed adaptive quadrature methods can be applied to dynamically adjust the bin width. Our iPC-entropy protocol can be applied to estimate $TS_{config}$ for molecular conformations obtained from enhanced sampling techniques or machine learning. However, these sampling techniques will perturb their conformational free energy landscape and result in bias computed $TS_{config}$, one need to reweigh the $TS_{config}$ to ensure fair comparison across different ligand.

**Circular Correlation**

Similar to calculation of circular covariance, we applied angle-aware arithmatic to compute ciruclar correlation coefficient (ρ).

$$\rho_{xy} = \frac{\sum_{i=1}^{n}(x_i-\bar{x})(y_i-\bar{y})}{\sqrt{\sum_{i=1}^{n}(x_i-\bar{x})^2}\sqrt{\sum_{i=1}^{n}(y_i-\bar{y})^2}}$$

$$\bar{x} = \arctan\left(\frac{sin(x_1)+sin(x_2)+\cdots+sin(x_n)}{cos(x_1)+\cos(x_2)+\cdots+cos(x_n)}\right)$$

$$x_i - \bar{x} = \arctan\left(\frac{\sin(x_i)\cos(\bar{x}) - \sin(\bar{x})\cos(x_i)}{\cos(x_i)\cos(\bar{x})+\sin(x_i)\sin(\bar{x})}\right)$$

$\rho_{xy} = \ Circular\ Pearson\ Correlation\ Coeficient,\ \overline{x} = mean\ of\ dihedral\ angles,$

$x_i\ and\ y_i = side\ chain\ dihedral\ angles$

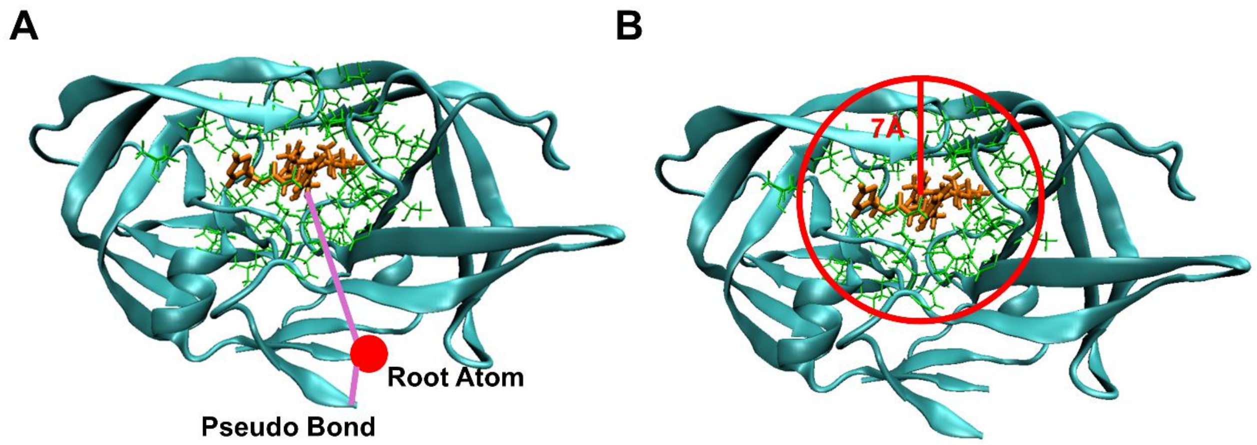

**Figure S1**. **RIT in complex with HIV Protease. (A)** The red dot indicates the root atoms. Two pseudo dihedral angles are included to correctly define the position of HIVp dimer and RIT. **(B)** 7Å from the ligand are defined as the flexible regions for configurational entropies calculation. 42 residues of the HIVp are selected based on the criteria. RIT is colored orange; The sidechains of 42 residues in the flexible regions are colored in green; The pseudo dihedral angles are colored in purple.

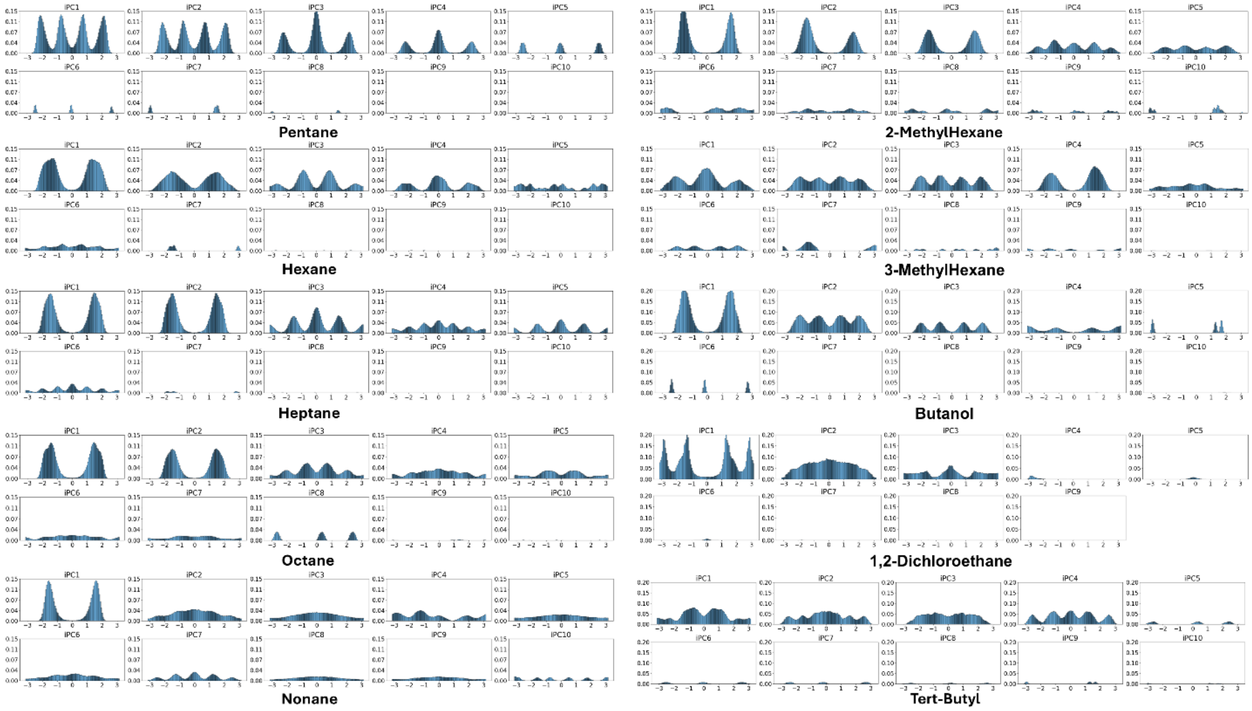

**Figure S2. Projecting the BAT internal coordinate on to corresponding eigenvectors of each iPC modes of organic molecules.** The PDFs demonstrate the molecular configurations of the first 10 iPCs and were used to compute $TS_{config}$.

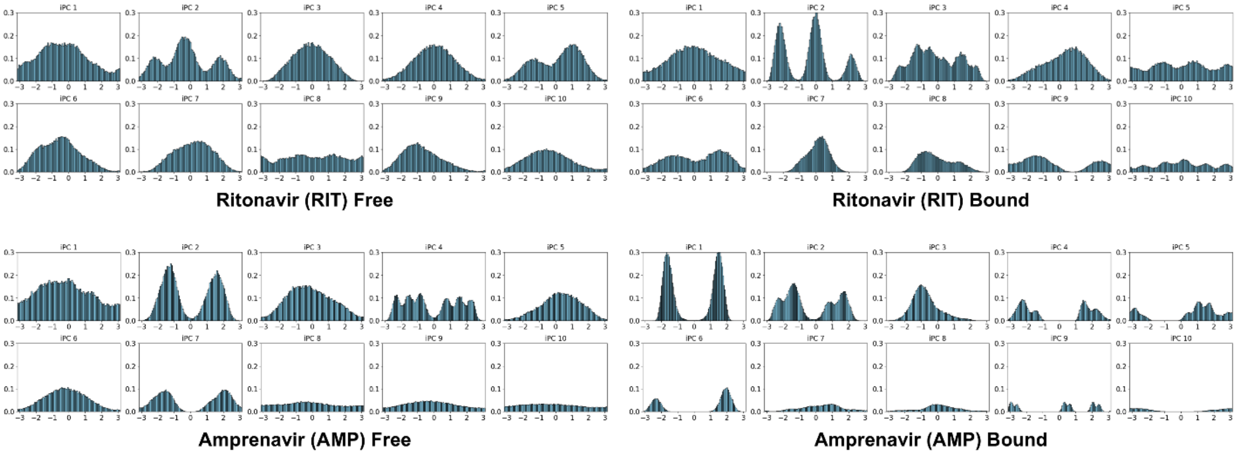

**Figure S3. Projecting the BAT internal coordinate on to corresponding eigenvectors of each iPC modes of RIT and AMP.** The PDFs of the first 10 iPC projections for free and bound state RIT and AMP.

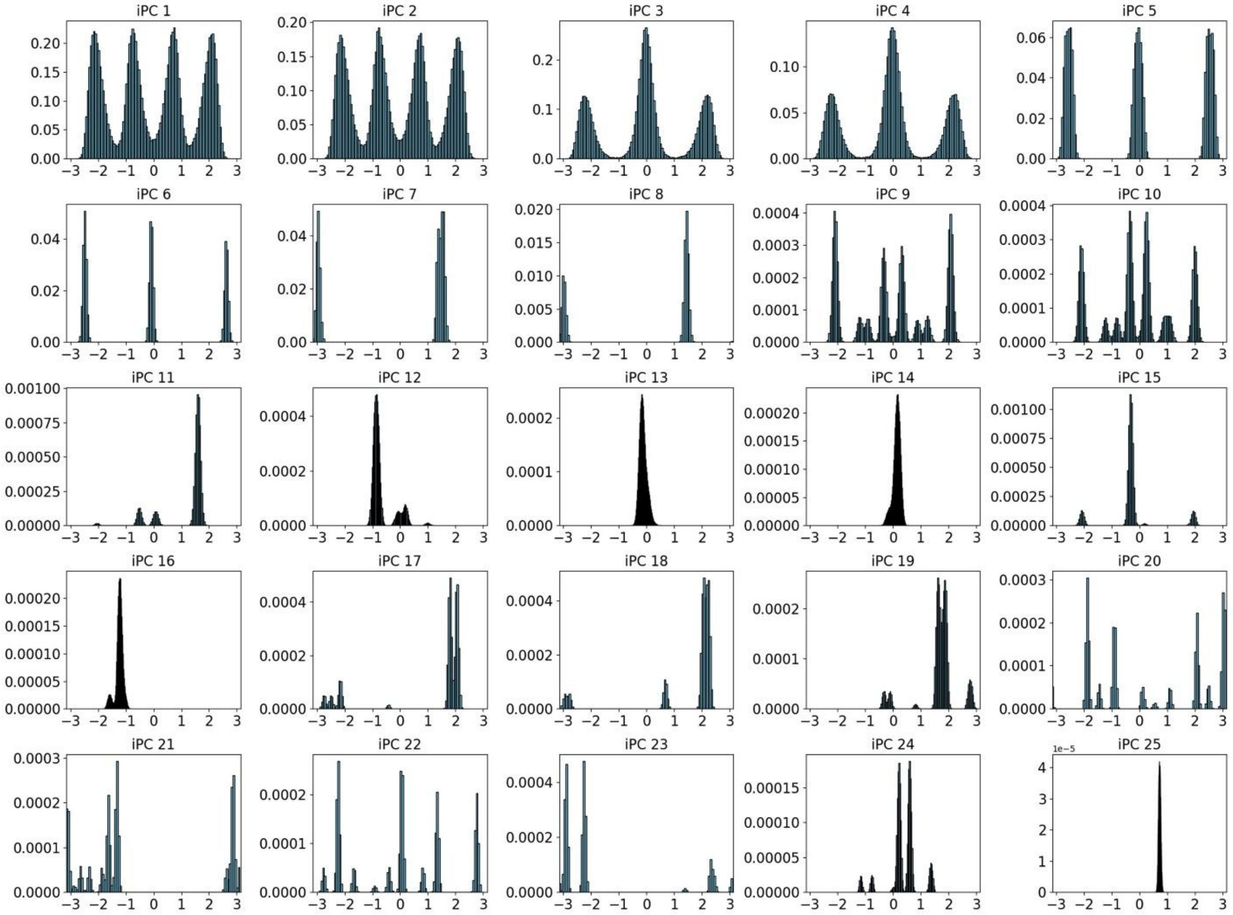

**Figure S4. De-coupling Molecular motions of pentane.** The PDFs of the projected data demonstrate the molecular configurations of the first iPCs showing both torsion angle rotations (first few PDFs) and bond/angle vibration (last few PDFs) where the last few PDFs shows Dirac delta function.

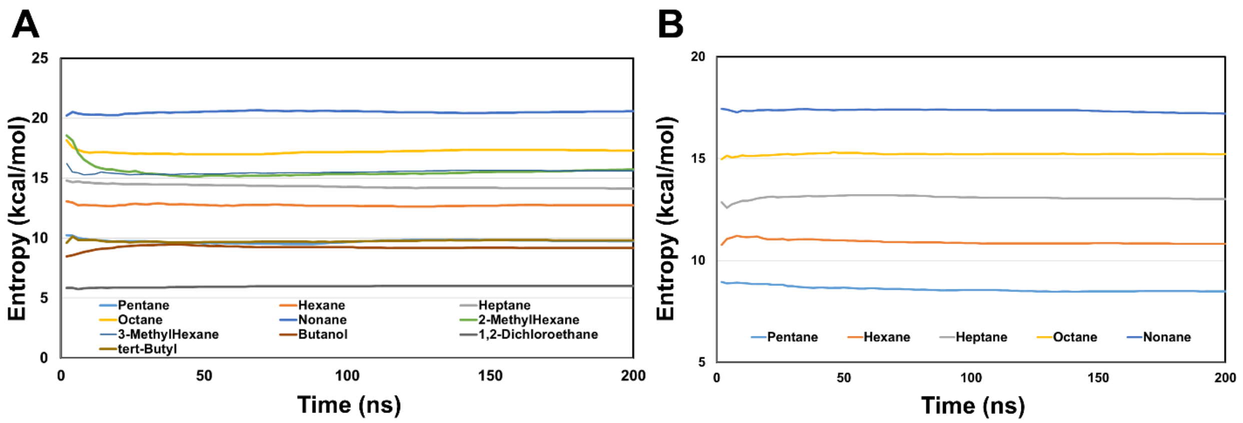

**Figure S5. Cumulative averaging of molar entropy, $TS_{config}$. (A)** Original Trajectories with GAFF2 force field, where the computed $TS_{config}$ all converge within 200 ns MD simulation length. **(B)** Angle restricted trajectories by increasing angle force constant to 300 pN yielded the same convergence within 200 ns MD simulation.

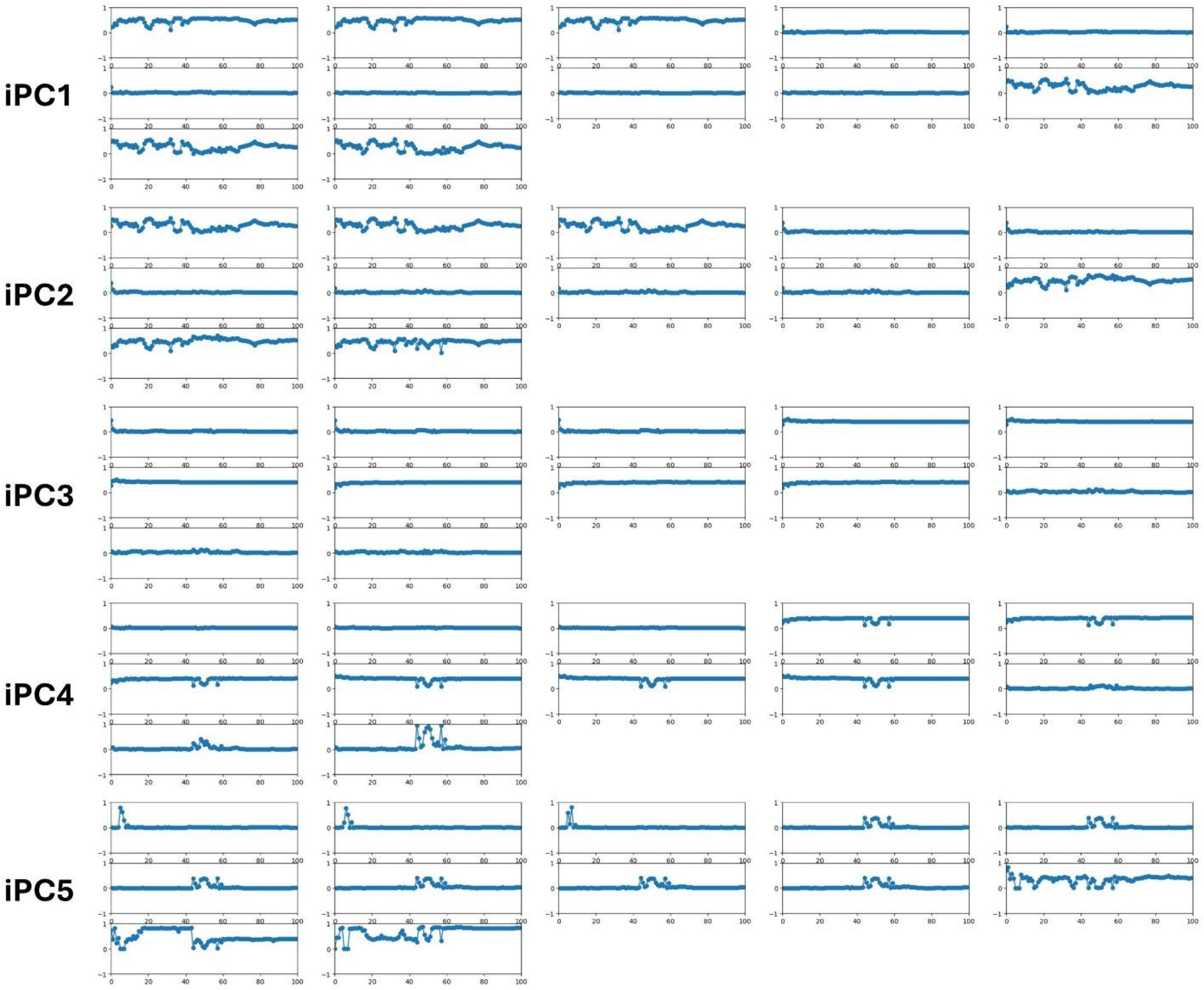

**Figure S6. Absolute values of the pentane eigenvectors over time of the first five iPC eigenmodes.** Eigenvectors are computed using 1-2k, 1-4k, 1-6k…1-200k frames of the MD trajectories which result in total of 100 points. The plot shows that eigenvectors stabilized after 80 (160k frames).

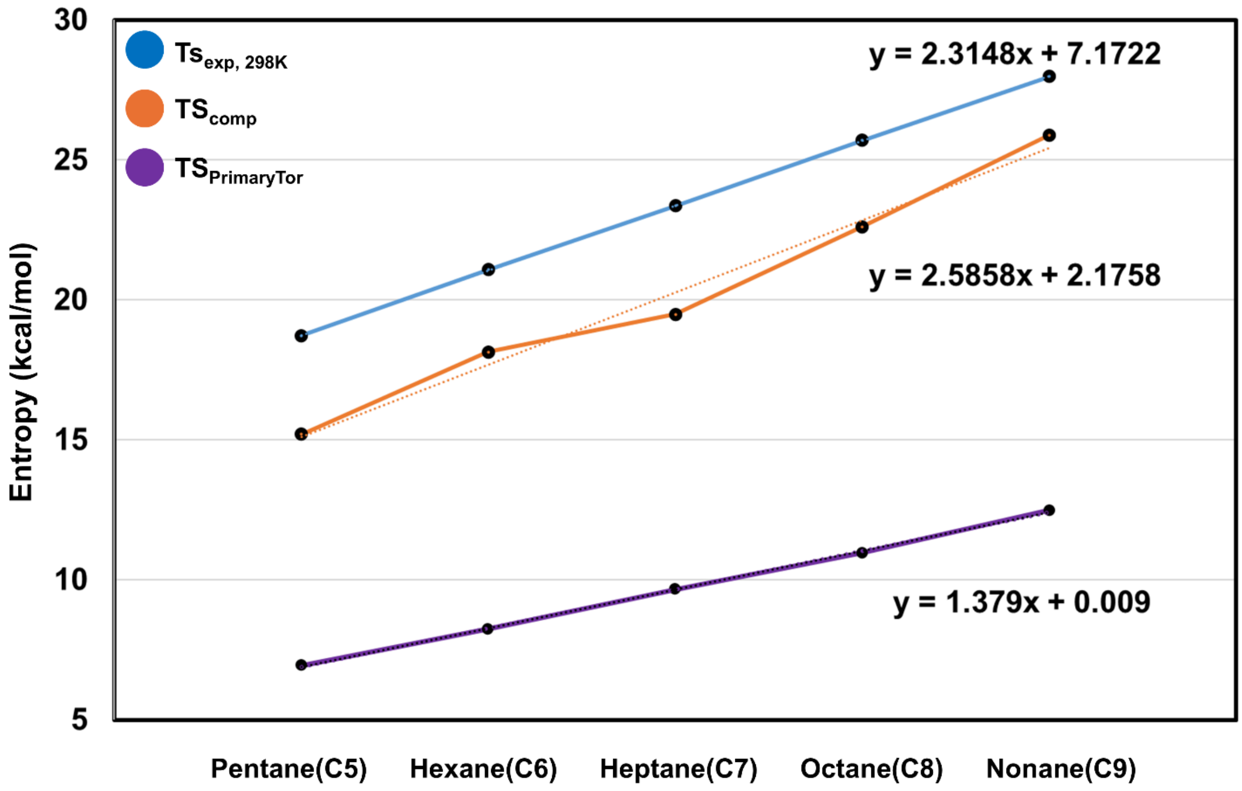

**Figure S7. Linear relationship between n-alkanes $C_5$ to $C_9$.** Blue shows experimental molar entropy ($TS_{exp,\ 298K}$). Red shows the computed molar entropy ($TS_{comp}$), and Purple shows the Torsion only entropy ($TS_{PrimaryTor}$) computed using all rotatable bond with T-analyst program.

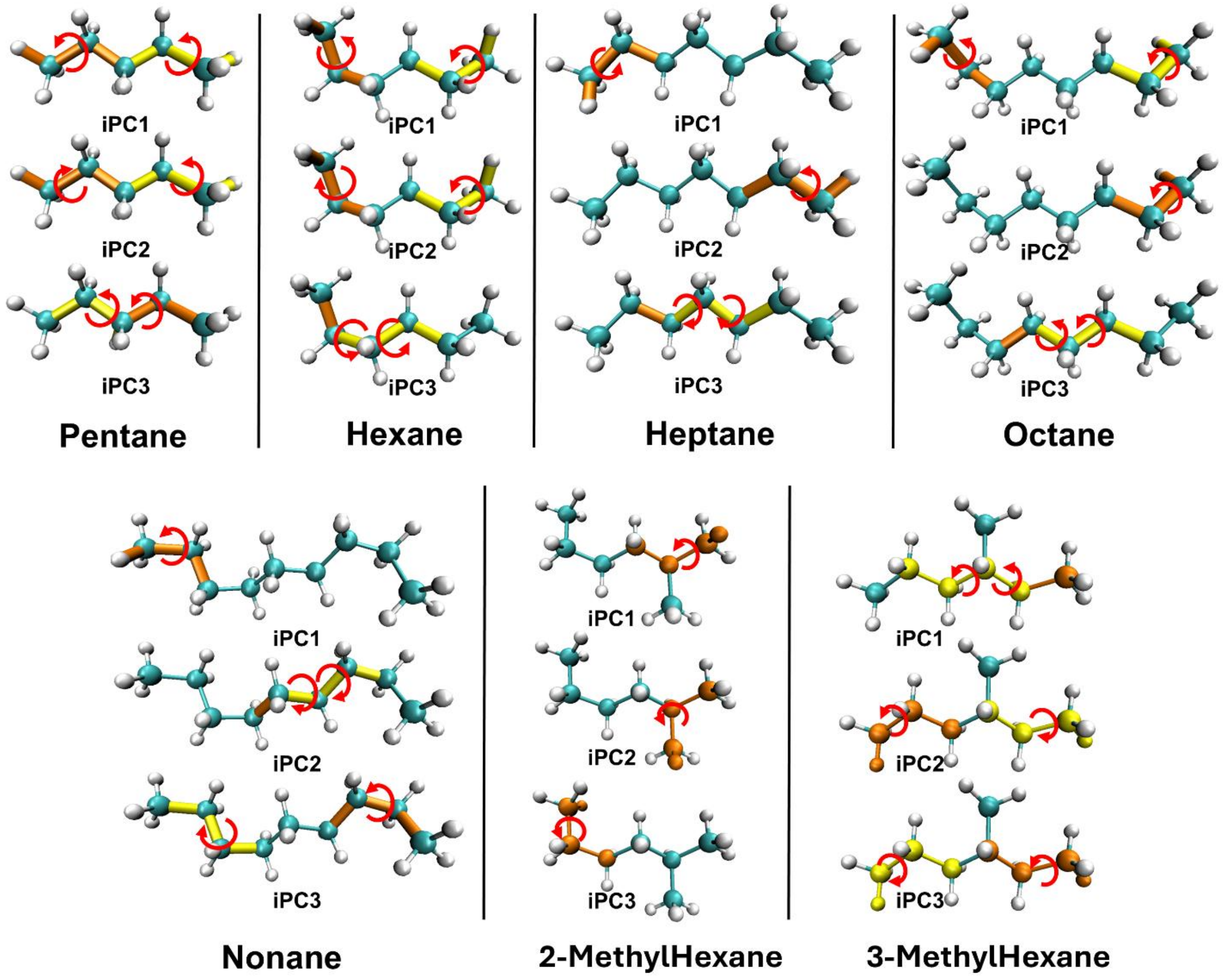

**Figure S8. Essential dihedral rotation of organic molecules.** We show the top two (orange and yellow stick) essential dihedral rotations of alkanes in the first three iPC eigenmodes. The essential motions were obtained directly by computing the eigenvectors of each eigenmodes. The red circular arrows indicate the rotation.

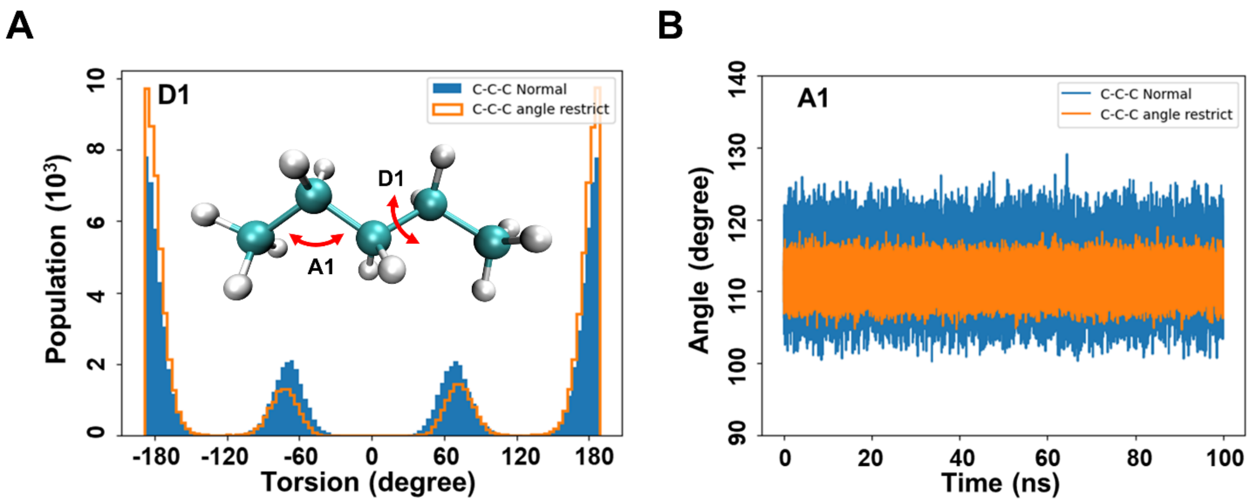

**Figure S9. Investigate the change in torsion rotation by comparing normal pentane simulation and angle-restricted pentane simulation. (A)** Probability distribution of dihedral angle, D1, of normal pentane and angle restricted pentane. **(B)** Angle vibration, A1, over the course of 100-ns MD simulation of normal pentane and angle restricted pentane.

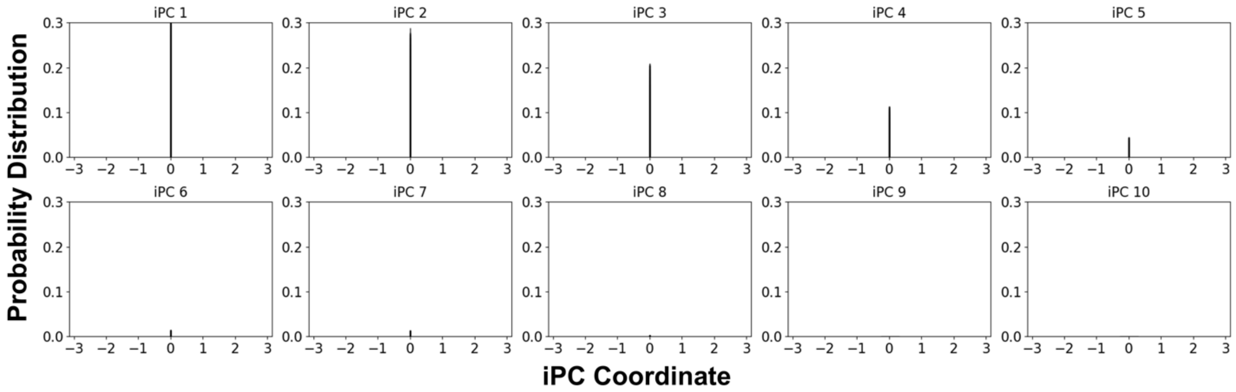

**Figure S10. Projection of pentane bond-angle bending on each eigenmodes.** Selecting only the eigenvectors of angle bending for projections. We observe only sharp peaks which contribute insignificantly to entropy calculation.

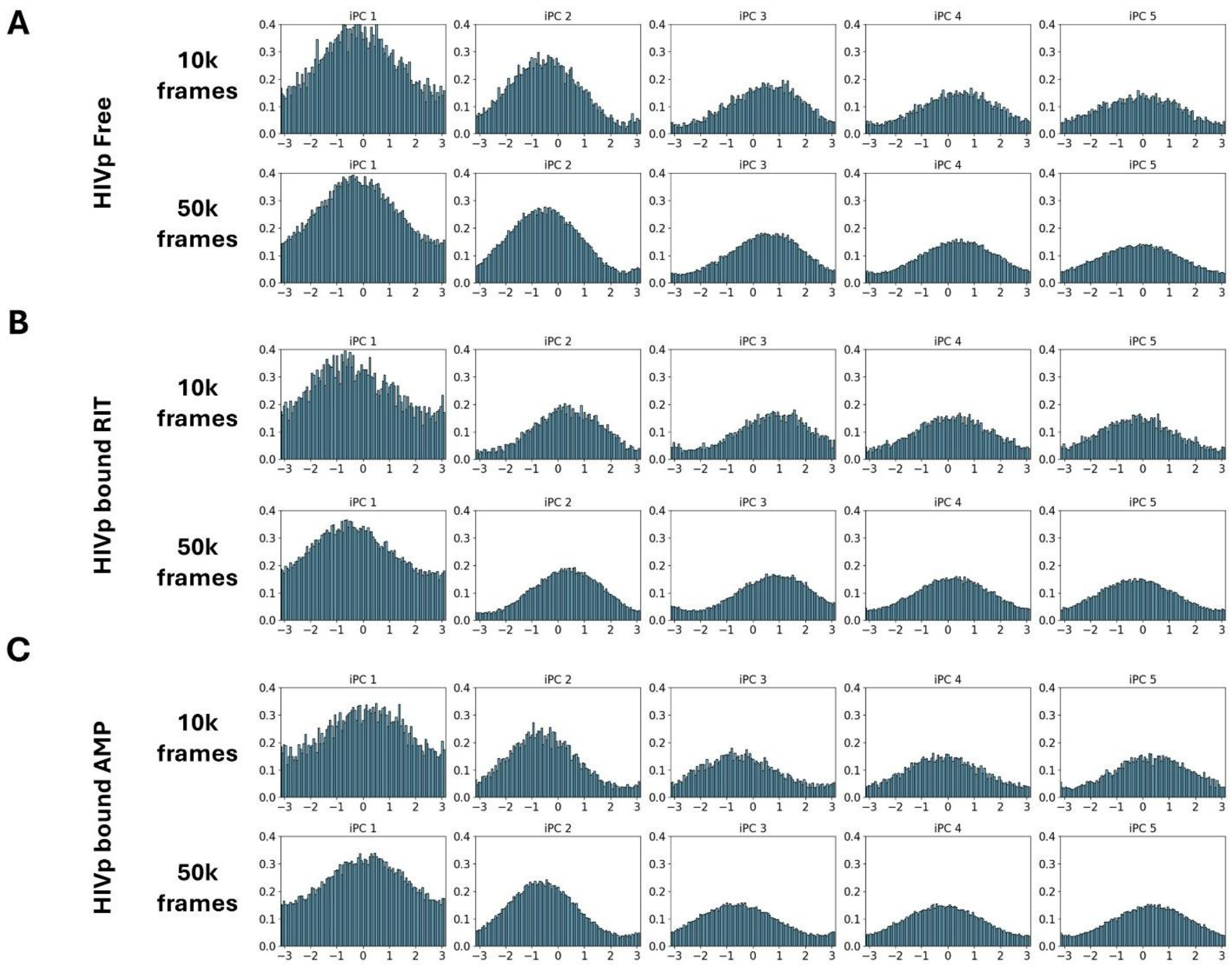

**Figure S11. Smoothness of the probability distribution function required for accurate numerical integration**. Compare 10,000 and 50,000 frames of the first 5 iPCA projection for **(A)** HIVp free state **(B)** HIVp bound with RIT **(C)** HIVp bound with AMP.

**Table S1.** Eigenvalues and Entropy (TS) of the first three iPC mode of n-alkanes $C_5$ to $C_9$.

| | | | | | | |
|---|---|---|---|---|---|---|
| | Pentane | | Hexane | | Heptane | |
| | Eigenvalues | TS | Eigenvalues | TS | Eigenvalues | TS |
| iPC1 | 9.97 | 2.22 | 9.42 | 2.11 | 8.98 | 1.96 |
| iPC2 | 8.17 | 2.11 | 6.93 | 2.10 | 8.81 | 1.94 |
| iPC3 | 5.63 | 1.71 | 5.90 | 2.01 | 6.23 | 2.00 |
| | Octane | | Nonane | | | |
| | Eigenvalues | TS | Eigenvalues | TS | | |
| iPC1 | 9.38 | 2.01 | 8.87 | 1.84 | | |
| iPC2 | 7.97 | 1.92 | 6.99 | 2.20 | | |
| iPC3 | 6.30 | 2.11 | 5.32 | 2.05 | | |

**Table S2.** Eigenvalues and Entropy (TS) of the first six iPC mode of free and bound RIT and AMP.

| | | | | |
|---|---|---|---|---|
| | Free RIT | | Bound RIT | |
| | Eigenvalues | TS | Eigenvalues | TS |
| iPC1 | 10.6 | 2.45 | 9.64 | 2.40 |
| iPC2 | 8.56 | 2.28 | 8.7 | 2.06 |
| iPC3 | 7.73 | 2.10 | 8.60 | 2.26 |
| iPC4 | 7.65 | 2.15 | 7.07 | 2.13 |
| iPC5 | 7.56 | 2.17 | 6.3 | 2.18 |
| iPC6 | 7.43 | 2.13 | 5.76 | 2.09 |
| | Free AMP | | Bound AMP | |
| | Eigenvalues | T | Eigenvalues | TS |
| iPC1 | 12.4 | 2.57 | 9.11 | 1.86 |
| iPC2 | 9.3 | 2.17 | 6.61 | 1.98 |
| iPC3 | 8.62 | 2.27 | 4.25 | 1.57 |
| iPC4 | 6.70 | 2.09 | 2.93 | 1.40 |
| iPC5 | 5.94 | 2.00 | 2.86 | 1.44 |
| iPC6 | 5.31 | 1.96 | 2.01 | 0.95 |

**Table S3.** Computed configurational entropy, $TS_{config}$ and $TS_{PrimaryTor}$.

| | $TS_{config}$ | $TS_{PrimaryTor}$ |
|---|---|---|
| Pentane | 9.76 | 6.95 |
| Hexane | 12.74 | 8.25 |
| Heptane | 14.13 | 9.66 |
| Octane | 17.29 | 10.96 |
| Nonane | 20.59 | 12.49 |

$TS_{config}$ is solute entropy with the Jacobians included. $TS_{PrimaryTor}$ was computed using T-analyst program which considered all rotatable bonds.

**Table S4.** Computed entropy $TS_{angle\text{-}restrained}$ upon drug RIT or AMP binding to HIVp in standard concentration (kcal/mol).

| | $TS_{angle\text{-}restrained}$ | $TS_{config}$ | $T\Delta S_{angle\text{-}restrained}$ | $T\Delta S_{config}$ | $T\Delta S^o_{angle\text{-}restrained}$ | $T\Delta S^o_{config}$ |
|---|---|---|---|---|---|---|
| Free RIT | 33.05 | 44.72 | | | -30.91 | -32.53 |
| Bound RIT | 19.23 | 24.22 | | | | |
| RIT | | | -13.82 | -20.50 | | |
| Free HIVp | 214.27 | 214.27 | | | | |
| RIT bound HIVp | 197.18 | 202.24 | | | | |
| HIVp | | | -17.09 | -12.03 | | |
| Free AMP | 22.62 | 29.67 | | | -27.44 | -25.70 |
| Bound AMP | 7.05 | 13.79 | | | | |
| AMP | | | -15.57 | -15.88 | | |
| Free HIVp | 214.27 | 214.27 | | | | |
| AMP bound HIVp | 202.40 | 204.45 | | | | |
| HIVp | | | -11.87 | -9.82 | | |

Changes of entropy of binding $T\Delta S_{angle\text{-}restrained}$ was computed using **equation 20**. Comparing $T\Delta S^o_{angle\text{-}restrained}$ and $T\Delta S^o_{config}$, we observed ~2kcal/mol difference.

**Table S5.** Comparing the entropy $TS_{config}$ when using 50K and 10K frames for numerical integration in **equation 20**.

| | $TS_{config}$ | |
|---|---|---|
| | 50K frames | 10K frames |
| Free RIT | 44.72 | 44.61 |
| Bound RIT | 24.22 | 24.20 |
| Free HIVp | 214.27 | 213.89 |
| RIT bound HIVp | 202.24 | 201.88 |
| Free AMP | 29.67 | 29.61 |
| Bound AMP | 13.79 | 13.75 |
| Free HIVp | 214.27 | 213.89 |
| AMP bound HIVp | 204.45 | 204.12 |
